# Alignment of spatial genomics and histology data using deep Gaussian processes

**DOI:** 10.1101/2022.01.10.475692

**Authors:** Andrew Jones, F. William Townes, Didong Li, Barbara E. Engelhardt

## Abstract

Spatially-resolved genomic technologies have allowed us to study the physical organization of cells and tissues, and promise an understanding of the local interactions between cells. However, it remains difficult to precisely align spatial observations across slices, samples, scales, individuals, and technologies. Here, we propose a probabilistic model that aligns a set of spatially-resolved genomics and histology slices onto a known or unknown common coordinate system into which the samples are aligned both spatially and in terms of the phenotypic readouts (e.g., gene or protein expression levels, cell density, open chromatin regions). Our method consists of a two-layer Gaussian process: the first layer maps the observed samples’ spatial locations into a common coordinate system, and the second layer maps from the common coordinate system to the observed readouts. Our approach also allows for slices to be mapped to a known template coordinate space if one exists. We show that our registration approach enables complex downstream spatially-aware analyses of spatial genomics data at multiple resolutions that are impossible or inaccurate with unaligned data, including an analysis of variance, differential expression across the *z*-axis, and association tests across multiple data modalities.

## 1 Introduction

Spatially-resolved genomic technologies hold the promise to understand the spatial organization, variation, and local effects of cellular morphology, somatic mutations, gene expression, protein expression, and other cellular phenotypes (Ståhl et al., 2016; Rodriques et al., 2019; Stickels et al., 2021; Lee et al., 2021; Zhao et al., 2021). However, it remains difficult to jointly analyze multiple phenotypic readouts from these technologies due to inevitable spatial warping and biological variation across slices, samples, and individuals. Moreover, the coordinate system, spatial resolution, slice, scale, field-of-view, and data modality of these data vary widely across different technologies, making it difficult to analyze multiple spatial genomics slices jointly.

Here, we present a probabilistic model that aligns the local coordinate systems of spatially-resolved genomics and histology datasets across tissue slices, individuals, and technologies. Our model is well-suited for spatially-resolved measurements of cellular morphology, somatic mutations, gene expression, protein expression, and other cellular phenotypes, each of which may be collected at sub-cellular, cellular, or super-cellular resolution. Given a set of misaligned slices, our approach is to iteratively estimate a robust common coordinate system — a set of spatial locations along with the distribution of phenotypic readout values found at each location — using these slices, and map the local coordinates from each slice onto the common coordinate system. Our model, which leverages similarities in both the spatial structure and phenotypic readouts between slices, enables the creation of a common coordinate system onto which heterogeneous slices may be mapped and then analyzed jointly. Using this framework, new slices may also be projected into a given coordinate system, of which there are only a few (Sunkin et al., 2012; Rozenblatt-Rosen et al., 2020).

Our proposed generative model uses two stacked Gaussian processes (GPs) to align spatial slices across technologies and samples in a two-dimensional, three-dimensional, or potentially spatiotemporal coordinate system. Given a location in a slice, the first layer of the fitted model maps this location to its corresponding location in the common coordinate system, and the second layer generates the distribution of phenotypic readout values at that location (e.g., the distribution of gene expression for each gene).

Our approach opens the door to the creation of large tissue and organ atlases with a shared coordinate system using collections of tissue samples. Our model allows for straight-forward downstream analyses on the aligned slices, including imputation, completion of sparse measurements, analysis of variation, spatially-resolved differential expression, mapping between data modalities, and uncertainty estimates for each of these analyses. Moreover, technologies with different phenotypic readout modalities can be jointly analyzed under our model, including spatial transcriptomics, spatial protein expression, and histology images such as hematoxylin and eosin (H&E) stains (Fischer et al., 2008).

### 1.1 Background

#### 1.1.1 Spatially-resolved genomics technologies

In the past decade, a number of technologies have been developed to obtain spatially-resolved molecular measurements in biological tissues. A common readout of these technologies is gene expression levels (quantification of RNA transcripts), where the spatial locations of those transcripts are collected simultaneously with gene expression measurements (Lubeck and Cai, 2012; Eng et al., 2019; Ståhl et al., 2016; Rodriques et al., 2019; Lee et al., 2021). In addition to gene expression, technologies have been developed to measure genotype and somatic mutations, protein expression, chromatin accessibility, and other molecular readouts with spatial resolution (Goltsev et al., 2018; Keren et al., 2019; Thornton et al., 2021; Zhao et al., 2021). These protocols also often collect matching histology images in the form of H&E stains. As new technologies have been developed, several computational models and analysis pipelines have been proposed for processing and downstream analyses of single slice data (Velten et al., 2020; Townes and Engelhardt, 2021; Atta and Fan, 2021; Verma and Engelhardt, 2020; Svensson et al., 2018; Dries et al., 2021; Palla et al., 2021).

Although these methods have allowed for new scientific discoveries, it remains difficult to analyze multi-slice datasets due to spatial distortions in the tissue sample that occur during preparation and tissue collection. Furthermore, the various spatial genomics platforms range widely in their fields-of-view, their levels of spatial resolution, and the number of phenotypic features they measure. The current standard analysis approach — in which each slice is analyzed in isolation — drastically reduces the statistical power of the analyses, or prohibits these analyses entirely. Thus, there remains a need for tools that enable a joint analysis across slices, samples, modalities, scales, and technologies.

#### 1.1.2 Registration and hyperalignment

The problem of integrating disparate spatial and temporal slices from multiple sources has arisen in several fields. The problem of spatial alignment has perhaps been most thoroughly studied in the context of functional magnetic resonance imaging (fMRI) data and is referred to as *registration* in that context (Brett et al., 2001; Klein et al., 2009; Lancaster et al., 2000).

At a given time point, an fMRI scan produces measurements on a three-dimensional grid across the brain across time, where the continuous level of blood flow is measured at each point (“voxel”) in the grid. Given multiple scans across days or subjects, the alignment problem is to warp the spatial coordinates of each voxel in a scan so that the matching brain regions for each patient are aligned and their (*x, y, z*) coordinate systems refer to approximately the same functional voxel in the brain.

Two major efforts to align fMRI scans have emerged: template-based registration and hyperalignment. Template-based registration methods seek to align scans from different subjects to a pre-defined template spatial coordinate system. This template is typically defined as a single subject’s scan or as the average of a collection of scans across multiple manually-aligned subjects. The most popular approach uses a “template brain” developed at the Montreal Neurological Institute (MNI) (Evans, 1992; Collins et al., 1994), which is the functional equivalent of a tissue atlas. Then, collected MRI and fMRI samples’ voxel coordinates are warped so that the new samples’ voxel coordinates match this template in terms of both location in the brain and voxel behavior across time.

Hyperalignment approaches seek to align different subjects’ data without a pre-defined template. In particular, hyperalignment methods compute scan-specific transformations of the voxel space such that each scan is aligned with the others, using the “centroid” of all scans as the template. Both linear (Haxby et al., 2011) and nonlinear (Lorbert and Ramadge, 2012) hyperalignment approaches have been developed. However, hyperalignment approaches are less concerned with creating an interpretable common coordinate system, instead focusing on estimating a generic latent representation of the scans.

More recently, a flexible nonparametric model called the Brain Kernel was developed for analysis of fMRI data (Wu et al., 2021). This model, similar to the model we propose here, is a two-layer deep Gaussian process (DGP), where the intermediate layer of hidden variables captures functional correlation between voxel locations across the brain. However, the Brain Kernel is not concerned with finding a common coordinate system for multiple fMRI samples; instead, the DGP estimates the observed spatial coordinates from one sample using a higher-dimensional (*>* 3) latent space that includes both (*x, y, z*) coordinates and additional latent coordinates in order to build a low-dimensional representation of the spatially-resolved covariance matrix of the voxel behavior.

Alignment methods developed for fMRI data are not easily extensible to the setting of spatial genomics and histology slices for a variety of reasons. First, curated structural and anatomical common coordinate systems are not available for the diversity of tissue types, developmental stages, and species that are studied using spatial genomics. Second, while the readout at each location for fMRI data is a scalar (a single number representing the level of blood flow), the readout in spatial genomics applications can be high dimensional, often having on the order of 10^3^ − 10^5^ sparse and noisy features. Finally, while the spatial resolution of fMRI scans tends to be one of a small number of values (a small set of standard resolutions for fMRI machines), there is a wide diversity of spatial technologies, each with their own resolution and field-of-view. Thus, there is a need for spatial genomics-specific alignment tools.

#### 1.1.3 Methods for spatial genomics slice alignment

Several computational methods have been proposed to align genomics RNA-sequencing data from multiple batches, labs, or time points (Butler et al., 2018; Hie et al., 2019; Stuart et al., 2019; Korsunsky et al., 2019).

In the spatial setting, to the best of our knowledge, just two approaches have been proposed for aligning samples’ spatial coordinates.

As one of the first efforts in this direction, Probabilistic Alignment of Spatial Transcriptomics Experiments (PASTE) (Zeira et al., 2021) was developed to align adjacent tissue slices in Spatial Transcriptomics (ST) data (Ståhl et al., 2016). PASTE uses an optimal transport framework to identify mappings between the spatial locations of adjacent slices. Its objective function trades off the transcriptional similarity and physical proximity of spatial locations. While PASTE is a robust and fast framework, it is limited to linear alignments, which are often insufficiently expressive for complex distortions of data. Furthermore, it does not have an associated generative model, precluding the quantification of uncertainty in the alignment.

A landmark-based method relying on Gaussian process regression called Effortless Generic GP Landmark Transfer” (Eggplant) was recently developed (Andersson et al., 2021). Eggplant projects the gene expression values of each misaligned slice onto a fixed template coordinate system. It requires the user to identify 𝓁 landmark locations on each misaligned sample and the template sample. While Eggplant can account for spatial correlation in gene expression levels, it performs this template transfer independently for each slice and each gene, ignoring any correlation structure between them. Moreover, the selection of landmark locations may be difficult for slices from tissues without a canonical structure, such as tumors, and requires annotating existing landmarks shared across slices.

### 1.2 Contributions of this paper

In this work, we develop Gaussian Process Spatial Alignment (GPSA), a Bayesian framework for spatial alignment and analysis of multiple spatially-resolved genomics and histology slices. The contributions of this paper are as follows.

- Given a set of spatial genomics tissue slices with distorted or unaligned local coordinate systems, GPSA estimates a common coordinate system as a collection of latent variables from this set of disparate spatially-resolved measurements and aligns new samples to this estimated common coordinate system;
- GPSA can be used with an existing common coordinate system or a set of landmarks;
- GPSA jointly aligns datasets to the common coordinate system from multiple phenotypic readout modalities, including spatial gene expression, spatial protein expression, spatial somatic DNA mutations, spatial open chromatin regions, and histology images;
- GPSA allows for alignment of slices that do not overlap, or are parallel slices of the same tissue, or are across individuals, or have different fields-of-view, scales, or resolutions;
- GPSA allows for imputation of missing data and creation of dense spatial readouts for an arbitrary collection of spatial coordinates within the coordinate system that can be used for downstream analysis;
- These common coordinate systems may be used for a variety of multi-slice downstream tasks. For example, one could use these aligned coordinates to identify spatially variable genes (Svensson et al., 2018), explore low-dimensional structure across multiple samples (Townes and Engelhardt, 2021; Velten et al., 2020), project nonspatial (disassociated) single-cell data onto the spatially-resolved landscape (Verma and Engelhardt, 2020), and create atlases of spatial genomics data for entire tissues or organisms within a common coordinate system (Hawrylycz et al., 2012).

## 2 Problem definition and notation

Here, we formalize the problem we wish to solve. We define a *spatially-resolved slice* or *slice* from a spatial genomics or histology technology as a set of pairs 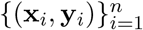, where **x**_*i*_ ∈ ℝ^*D*^ is a vector of spatial coordinates encoding a single slice’s relative location in a *D*-dimensional space, and **y**_*i*_ ∈ R^*p*^ is a vector of measured readout features at this location. Typically, *D* ∈ {2, 3, 4} in biomedical applications, where *D* = 4 corresponds to the spatiotemporal setting. We focus on *D* = {2, 3} in this paper. Following convention, we refer to a single location **x**_*i*_ as a *spot*, which may refer to a single cell, a subcellular location, a collection of cells, or a single pixel depending on the technology. We arrange the observations from a slice into two data matrices: the spots’ relative locations **X** ∈ ℝ^*n×D*^ and the phenotypic readouts associated with those spots **Y** ∈ ℝ^*n×p*^.

To give concrete examples, in *spatial transcriptomics* applications, **x**_*i*_ encodes a spatial location on a single tissue slice, and **y**_*i*_ is a vector of RNA transcript counts at this location for each of *p* genes. In a *histology* setting, **x**_*i*_ is a pixel location, and **y**_*i*_ is a vector containing the *p* image color channel readouts.

We assume we have *S* spatially-resolved slices collected from the same tissue type and similar tissue region. Often, these slices will be adjacent slices from a single tissue, but as we will show in our results, our approach is extensible to datasets collected from different tissue samples or individuals. Suppose slice *s* (*s* ∈ {1, …, *S*}) contains *n*_*s*_ spots, and let 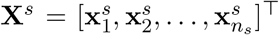 denote its spatial locations. Similarly, let 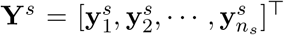 be the *s*th readout of feature values. We denote the total number of spots across slices as 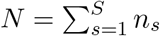. We note that, in our framework, the slices may have different total numbers of spots, and may be on different scales.

Our goal is to align these *S* slices’ spatial coordinates by creating a common coordinate system such that the matching anatomical, structural, and functional regions of each slice are mapped to the same absolute locations in the common coordinate system. To do this, we seek correspondences between both the spatial coordinates and phenotypic readouts of each slice. Specifically, we seek *S* vector-valued warping functions *g*^1^, *g*^2^, …, *g*^*S*^, with *g*^*s*^ : ℝ^*D*^ → ℝ^*D*^, each of which maps a slice’s observed relative spatial coordinates into a shared common coordinate system. Let 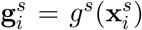 denote the evaluation of the *s*th warping function at spatial location 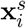. We call 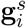 the *aligned spatial location* of this spot, and let the full set of aligned spatial locations be denoted as 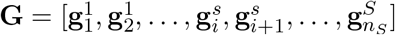.

Our goal is to estimate these warping functions 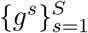 such that any two samples mapped to nearby points in the common coordinate system, 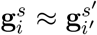, are structurally and functionally similar to one another. We consider three approaches, that show the powerful behavior of our probabilistic model under uncertainty and censored information. First, we treat the multiple slices as biological replicates to leverage both spatial information and the measured readouts for alignment. Second, we consider multiple data modalities of the same biological system, assuming the data come from approximately the same location in the absolute coordinate system to leverage the spatial locations and all modalities jointly. Third, we use multiple slices and infer their relationship along an unobserved *z*-axis, assuming that the measured readouts vary across the *z*-axis in a smooth way. The flexibility of our Gaussian process framework allows each of these three approaches to alignment.

## 3 Methods

### 3.1 Gaussian Process Spatial Alignment (GPSA)

Here, we propose Gaussian Process Spatial Alignment (GPSA), a Bayesian model for aligning spatial genomics and histology slices whose spatial coordinates are distorted or on different systems across the slices. GPSA consists of a two-layer deep Gaussian process (DGP, Damianou and Lawrence (2013)), where the first layer GP maps the observed spatial coordinates of a slice onto a latent common coordinate system, and the second layer maps the common coordinates onto the observed phenotypic readout values. We describe each of these layers in turn.

#### First layer: Warping functions

GPSA places GP priors on the warping functions *g*^1^, …, *g*^*S*^ that map the observed spatial coordinates onto a common coordinate system. Focusing on the case with *D* = 2 spatial dimensions for demonstration, GPSA assumes

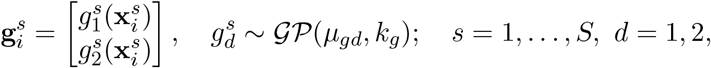

where 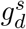 is the warping function for slice *s* whose output is the *d*th spatial dimension, *μ*_*gd*_ : ℝ^*D*^ → ℝ is a mean function, and *k*_*g*_ : ℝ^*D*^ × ℝ^*D*^ → ℝ is a positive definite covariance function. We specify the mean of the aligned spatial location to be equal to the observed location, *μ*_*gd*_(**x**) = *x*_*d*_, which encourages the aligned coordinate for a given spatial location to be centered around the observed location. This assumption is useful to avoid extreme warps that drastically shift the mean of each observed location.

#### Second layer: Modeling phenotypic readouts

GPSA then posits another set of functions 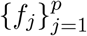 that describe the spatial organization of each phenotypic readout or feature (e.g., gene expression values) within the common coordinate system. We place a GP prior on these functions as well. Letting 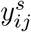 denote the value for feature *j* in spot *i* from slice *s*, GPSA assumes

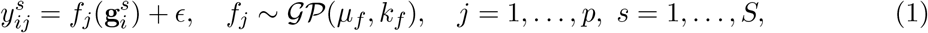

where ϵ ∼ 𝒩 (0, *∼*^2^) is a noise term, *μ*_*f*_ : ℝ^*D*^ → ℝ is a mean function, and *k*_*f*_ : ℝ^*D*^ ×ℝ^*D*^ → ℝ is a positive definite covariance function. Let 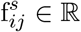 be the evaluation of *f*_*j*_ at input 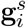. We specify *μ*_*f*_ = 0, as we assume the phenotypic readouts have been centered. Furthermore, let 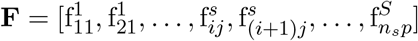 denote the full set of function evaluations.

The above model results in a two-layer DGP where, for each slice *s*, the DGP is made up of a composition of two functions, *f* ∘ *g*^*s*^. By placing GP priors on each of these functions, our model results in a two-layer DGP.

Whereas each slice has its own warping function *g*^*s*^, the functions modeling the spatial organization of phenotypic readouts in the common coordinate system, 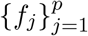, are shared across the slices. In other words, the first layer warps the locations of each spot in each slice to the common coordinate system while the second layer describes, at each location in the common coordinate system, the distribution of the readouts at that location captured from the aligned slices. Inference is thus guided by two competing objectives: retain the current relative position of each spot in a slice while warping each spot to ensure the readouts match as closely as possible with the distribution of readouts localized in the common coordinate system.

### 3.2 Multi-modality and low-rank modeling of the phenotypic readouts

Our GPSA allows for joint modeling of multiple types of readout modalities. For example, many experiments collect both spatial gene expression profiles and histology images for each tissue slice (Ståhl et al., 2016; Vickovic et al., 2019). These modalities contain complementary information, and it is of interest to analyze both modalities across multiple slices jointly in a common coordinate system. To do this, we augment our model of the phenotypic readouts (Equation 1) to include a separate likelihood for each modality, allowing for straightforward multi-modality alignment. See Appendix 6.3.6 for more details.

Although the measurements at each location are typically high-dimensional, the readout features tend to be highly correlated with one another, and this correlation is often well described (based on scRNA-seq experiments) by a lower-dimensional (*< p*) manifold (Linderman, 2021; Zeisel et al., 2015; Pierson and Yau, 2015; Ding et al., 2018). To account for this, GPSA allows for low-rank modeling of the outcome variables through a linear model of coregionalization (LMC; Goulard and Voltz (1992); Appendix 6.3.5), which models the outcomes as a weighted linear combination of a small number of GPs.

### 3.3 Posterior inference for the common coordinate system

We have two statistical objectives with the GPSA model: estimating the common coordinate system, as represented by the latent variable **G**, and estimating the warped, denoised, and localized values for the phenotypic readouts for each slice, represented by **F**. The common coordinate system gives us an atlas of the system; the warped and smoothed readouts may be used for downstream analysis of the aligned slices. Thus, in our Bayesian GPSA framework, the posterior distribution of interest is

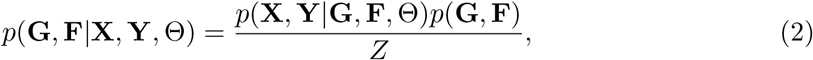

where the vector Θ contains the parameters for the mean and covariance functions and *Z* = *p*(**X, Y**|Θ) is a normalizing constant. However, *Z* is analytically intractable in DGPs (Damianou and Lawrence, 2013). Thus, we use stochastic variational inference with inducing variables to approximate the posterior distribution over **G** and **F** ((Hensman et al., 2013), see Appendix 6.3.3).

### 3.4 *De novo* and *template-based* common coordinate systems

Using GPSA, we propose two ways to align a set of slices: *de novo* alignment and *template-based* alignment. A *de novo* alignment estimates a common coordinate system from scratch using the slices while simultaneously projecting these slices into the common coordinate system. Alternatively, if a common coordinate system exists for a tissue (and organism, developmental phase, disease status, etc.) of interest, a *template-based* alignment maps the samples to this given coordinate system. This is accomplished by fixing the pre-defined common coordinate system as a non-warped slice within the set of aligned coordinates 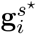, or equivalently fixing the warping function of the template slice to the identity, 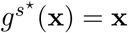. In practice, to avoid extreme warps in *de novo* alignment, we recommend arbitrarily choosing one of the input samples to fix as the canonical coordinate system.

## 4 Experiments

### 4.1 Model settings and preprocessing for experiments

In our experiments, which are restricted to slices that approximately share fields-of-view, we normalize all spatial coordinates so that the minimum *x* and *y* coordinate values are 0, and the maximum coordinate values are 10.

For all experiments, we specify the mean function of the GP prior for the warping functions to be the identity function. This choice is motivated by our expectation that most distortions in tissue samples will be relatively small and local, with large translations between slices being uncommon. We use the radial basis function (RBF) covariance function for the first layer GPs. The RBF covariance function is given by

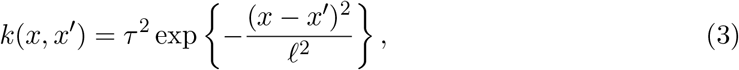

where 𝓁 is the length scale parameter, and *τ* ^2^ is the spatial variance parameter. For the second layer multi-output GP, with an LMC covariance function, we estimate the covariance function parameters using maximum likelihood as part of our inference procedure. For the first layer GP (the warp GP), we fix the covariance function parameters before model fitting. Specifically, we fix the length scale as 𝓁 = 10 and the spatial variance or amplitude parameters as *σ*^2^ = 1 to ensure smooth and minimal warps. We found that these choices are relatively robust within a range.

### 4.2 Simulations

We first validate the accuracy and robustness of our model using synthetic data generated under a variety of settings.

#### 4.2.1 Recovery of true latent common coordinates

First, we generated synthetic spatial gene expression data for two slices from a known underlying common coordinate system in order to study the behavior of GPSA.

We began with a simple setting with a one-dimensional spatial coordinate system in order to visualize the behavior of GPSA. We sampled a set of unobserved spatial coordinates for *n* = 100 spatial locations in the interval [0, 10]. We then generated observed spatial coordinates for *S* = 2 slices by applying a GP warp (Equation 10). We sampled synthetic gene expression *y*_*ij*_ for sample *i* and gene *j* using a GP:

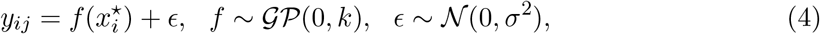

where 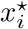 is the *i*th sample of the latent coordinates. We set *k* to be the RBF covariance function (Equation 3). In this experiment, we set *τ* ^2^ = 0.1. We fit GPSA to this dataset using a *de novo* alignment and extracted the aligned coordinates for each slice. We found that the warped coordinates were well aligned between the two slices and that the relationship between spatial coordinates and gene expression was preserved from the latent coordinates (Figure 1a). The mean squared error (MSE) for the aligned coordinates was 0.000134 (where an MSE of 0 indicates perfect performance), while the MSE of the original spatial coordinates was 0.0345. This result suggests that GPSA is able to align distorted and disparate samples accurately.

**Figure 1:**
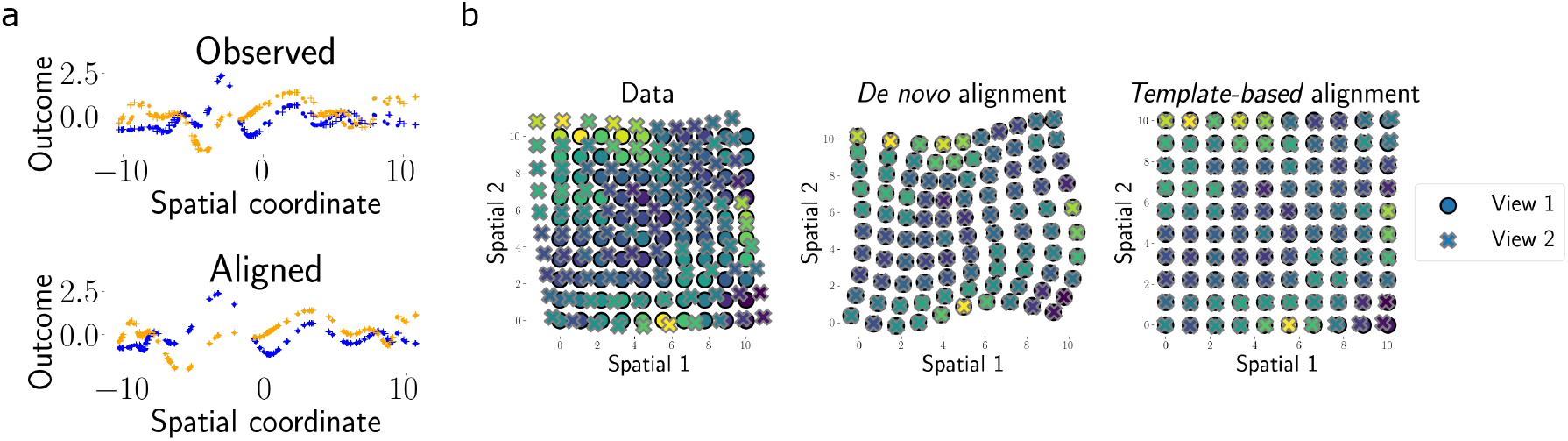
Demonstration of GPSA with synthetic data. (a) GPSA applied to a one-dimensional spatial coordinate system with *p* = 2 readout features (blue and orange) and *S* = 2 slices (dots and crosses). The x-axis shows the spatial coordinate of each sample, and the y-axis shows the readout values. The top panel shows the observed data, and the bottom panel shows the aligned coordinate system from GPSA. (b) GPSA applied to a one-dimensional spatial coordinate system with *p* = 10 readout features and *S* = 2 slices (dots and crosses). The x- and y-axes show the spatial coordinates of each sample. Points are colored by one feature value. The left panel shows the observed data, and the right panel shows the aligned coordinate system from GPSA.

Next, we extended this experiment to a more realistic setting in which the spatial coordinates are two dimensional. In this case, the unobserved coordinate system was a 15 × 15 grid containing *n* = 225 spatial locations. The observed spatial coordinates for the first slice were kept at the original, intact grid, and the observed coordinates for the second slice were generated by randomly warping the true coordinate system with the GP warp (Equation 10) using an isotropic RBF covariance function with length scale 𝓁 = 10 and spatial variance *τ* ^2^ = 0.5. We sampled synthetic gene expression values using the same approach as in Equation 4 from a GP with the true spatial coordinates as inputs. We fit the model twice this dataset: once using a *de novo* alignment and once using a *template-based* alignment with the first slice as the template. For each, we extracted the estimated warped coordinates of both slices. For both types of alignments, we again found that the coordinates were aligned between slices with minimal distortion using the latent common coordinate system (Figure 1b). The mean squared error (MSE) for the *de novo* and *template-based* aligned coordinates was 0.000537 and 0.00725, respectively, while the MSE of the original spatial coordinates was 0.733. These results indicate that GPSA is a viable model for aligning spatially-resolved slices whose spatial coordinates have been distorted using a latent shared coordinate system.

#### 4.2.2 Robustness of GPSA to observation noise

We next tested GPSA’s robustness to changes in the data generating process for the synthetic data. Specifically, we examined GPSA’s alignments for datasets generated under different *conditions*. Specifically, the conditions we varied were the number of readout features, the magnitude of distortion between the slices’ coordinate systems, and the level of noise variance in the readouts. To do so, we generated synthetic datasets with two-dimensional spatial coordinates using the same approach as in the previous section. We varied the number of readout features (“genes”) such that *p* ∈ {1, 20, 50} and generated datasets for each of these values. To vary the magnitude of the distortion between slices, we applied a GP warp (Equation 10) to the common coordinate system while varying the spatial variance *τ* ^2^ of the RBF kernel (Equation 3), which corresponds to a larger distortion between slices. We fit GPSA to each of these datasets using a *template-based* alignment with the first slice as the template, repeating the experiment five times for each condition. For comparison, we ran PASTE (Zeira et al., 2021) and extracted the aligned coordinates using the estimated linear transformation from PASTE. For each method, we computed the error of the aligned coordinates. Every pair of simulated slices contains spots at identical locations. We measured the error between the warped locations for each of these spots between every pair of slices, where a perfect warp of all slices would yield zero error:

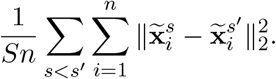

In this simulation experiment, we found that GPSA’s alignment error declined with more readout features and increased with greater distortion (Figure 2). GPSA achieved a substantially lower error than PASTE in all settings. We observe that this difference in error is largely due to the fact that PASTE only applies an linear transformation to each sample and is unable to account for local distortions. These results imply that our model is robust to nonlinear warpings, different magnitude distortions, and differences in the number of readout features.

**Figure 2:**
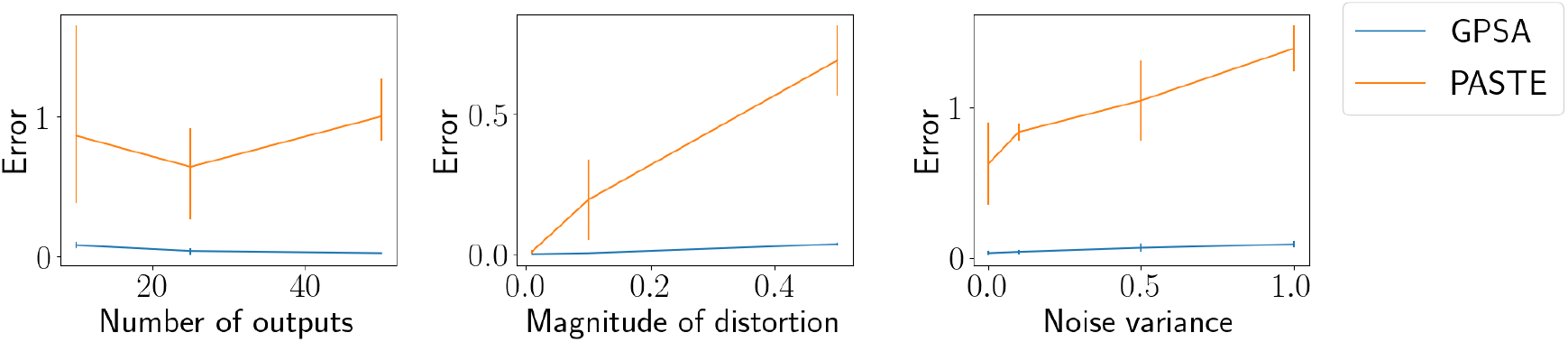
Benchmarking alignment error with synthetic data. (a) Alignment error (as measured by MSE between all pairwise aligned slices with shared original coordinates) of GPSA and PASTE (Zeira et al., 2021) across varying numbers of readout features. (b) Similar to (a), but shows the error across varying levels of distortion within the slices. (c) Similar to (a) and (b), but shows the error across different levels of variance in the synthetic expression data.

#### 4.2.3 Assessing alignment via readout prediction

Our experiments so far have tested whether GPSA can align the spatial coordinates of distorted samples. However, we expect that similar expression patterns across aligned slices should co-localize within the estimated common coordinate system. To test this, we attempted to predict held-out readout values using the posterior estimates of expression values localized within the common coordinate system. In particular, we repeated the two-dimensional experiment in Section 4.2.1, but we held out 20% of the samples from one of the slices. We then fit *template-based* GPSA and used the posterior estimates of expression values within the common coordinate system to predict the readout values at the held-out locations. Importantly, these predictions leverage information from all slices. We sampled from the variational posterior predictive distribution at each held-out location:

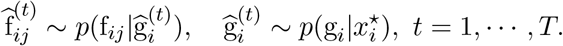

We then compared these predictions To make predictions, because the DGP posterior mean is not available in closed form, we approximate the posterior mean by drawing *T* = 10 samples for each location and taking the average of these as a point prediction, 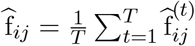. We then compared these predictions to the true readout values by computing the mean squared error. We compared GPSA to two baseline approaches: one approach that fits a GP to each slice separately (“separate” GP), and another approach that fits a GP to a naive concatenation of the original slices (“union” GP).

We find that GPSA achieves lower prediction error compared to the two baseline methods (Figure 3). This result suggests that GPSA estimates phenotypic readout value distributions localized within a common coordinate system that generalize in their predictive ability to held-out spots.

**Figure 3:**
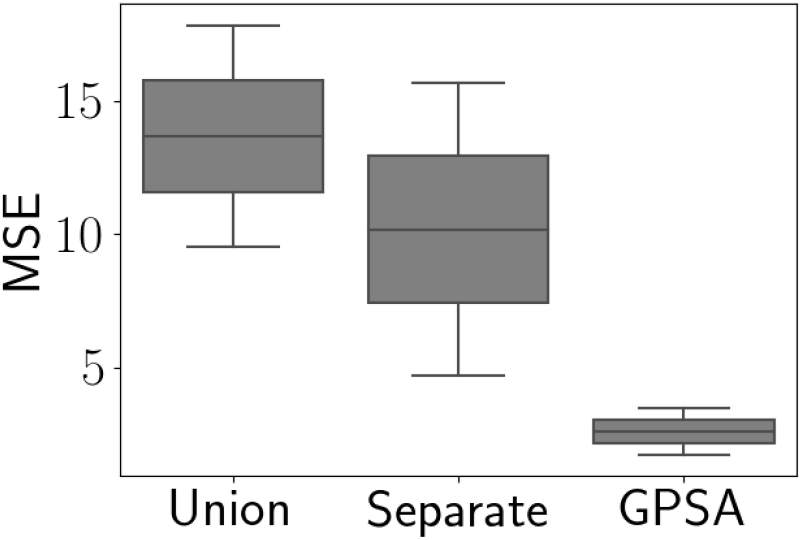
Prediction experiment with synthetic data. Mean squared error (MSE) for predictions of readout values versus ground truth on held-out spots from an aligned slice. “Union” represents predictions from a GP fit to a naive concatenation of the observed samples; “Separate” refers to predictions from independent GPs fit to each sample separately; and “GPSA” refers to predictions from GPSA fit across all samples using a latent common coordinate system.

#### 4.2.4 Aligning samples with different fields-of-view

In practice, the overlap in spatial coverage of samples may not be perfect. There may be settings in which we wish to align two spatial slices whose fields-of-views are different. To simulate this scenario, we generated data in which one slice was generated from a grid of spatial coordinates as before, and a second slice that was a small window fully contained within the original spatial coordinates and on the same spot grid (Figure 4a). We then fit *template-based* GPSA, using the larger slice as the template, to these two slices and extracted the aligned coordinates. For comparison, we also ran PASTE and compared results with those from GPSA.

**Figure 4:**
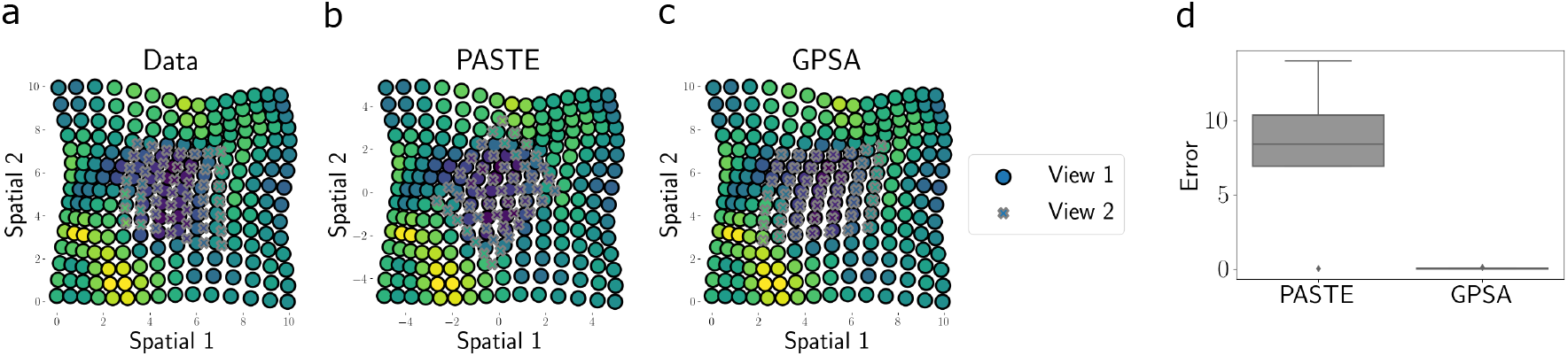
Aligning synthetic samples with different fields-of-view. (a) Synthetic data for two slices, one with a smaller field-of-view than the other. (b) Aligned coordinates from PASTE. (c) Aligned coordinates from GPSA. (d) Error for GPSA and PASTE aligned coordinates across five repetitions.

We found that GPSA is able to recover the alignment between two slices even when the fields-of-view are different (Figure 4a-c). As with the experiment above, the results from PASTE show substantially worse performance (Figure 4d).

### 4.3 Estimating common coordinate systems for spatial transcriptomics

Having validated GPSA as a viable model for robust spatial alignment of high-dimensional observations, we next applied GPSA to spatially-resolved genomics data. Below, we present analyses of data collected from three technologies: Spatial Transcriptomics (Ståhl et al., 2016), the Visium platform (10x Genomics, 2020), and Slide-seqV2 (Stickels et al., 2021). We also perform analyses with a set of images of H&E stains jointly with the spatial genomics data. An overview of the datasets used in this paper can be found in Table 1.

**Table 1.**
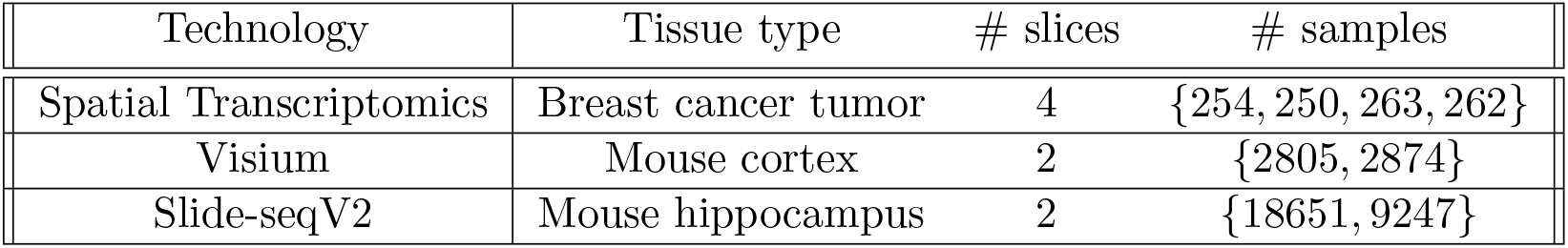
Spatial genomics datasets used in this paper.

| Technology | Tissue type | # slices | # samples |
| --- | --- | --- | --- |
| Spatial Transcriptomics | Breast cancer tumor | 4 | {254, 250, 263, 262} |
| Visium | Mouse cortex | 2 | {2805, 2874} |
| Slide-seqV2 | Mouse hippocampus | 2 | {18651, 9247} |

For all datasets, we performed preprocessing on these spatial sequencing data that includes removing mitochondrial genes; removing spatial locations with low counts; normalizing the readout at each spatial location by the total number of counts at that location; and log transforming, centering, and standardizing the gene counts. We further filter the datasets to include genes with spatial variability using a nearest-neighbor approach (see Appendix 6.3.8 for details). The spatial locations for each slice were normalized such that both coordinates were in the interval [0, 10], as the data were all produced with approximately the same field-of-view.

#### 4.3.1 Aligning Spatial Transcriptomics profiles of breast cancer samples

We tested our model on a dataset from the Spatial Transcriptomics (ST) technology (Ståhl et al., 2016) made up of four slices of a breast cancer tumor (Supplementary Figure 1).

We first sought to validate GPSA on the ST data by perturbing the samples with an artificial warp, and examining whether the common coordinate system estimated by GPSA approximately removed the perturbation. For this experiment, we analyzed each of the four samples separately. We applied a synthetic GP warp (Equation 10) with *τ* ^2^ = 0.5, 𝓁^2^ = 10 to each of the slices, and ran *de novo* GPSA on these misaligned samples. For comparison, we also ran PASTE, and we visualized the aligned coordinates for each of these methods.

We found that GPSA was able to recover the underlying coordinate system (Figure 5). Moreover, we observed that GPSA was able to correct the local spatial distortions in the observed spatial coordinates. On the other hand, we found that PASTE’s global correction was not sufficient to remove these distortions.

**Figure 5:**
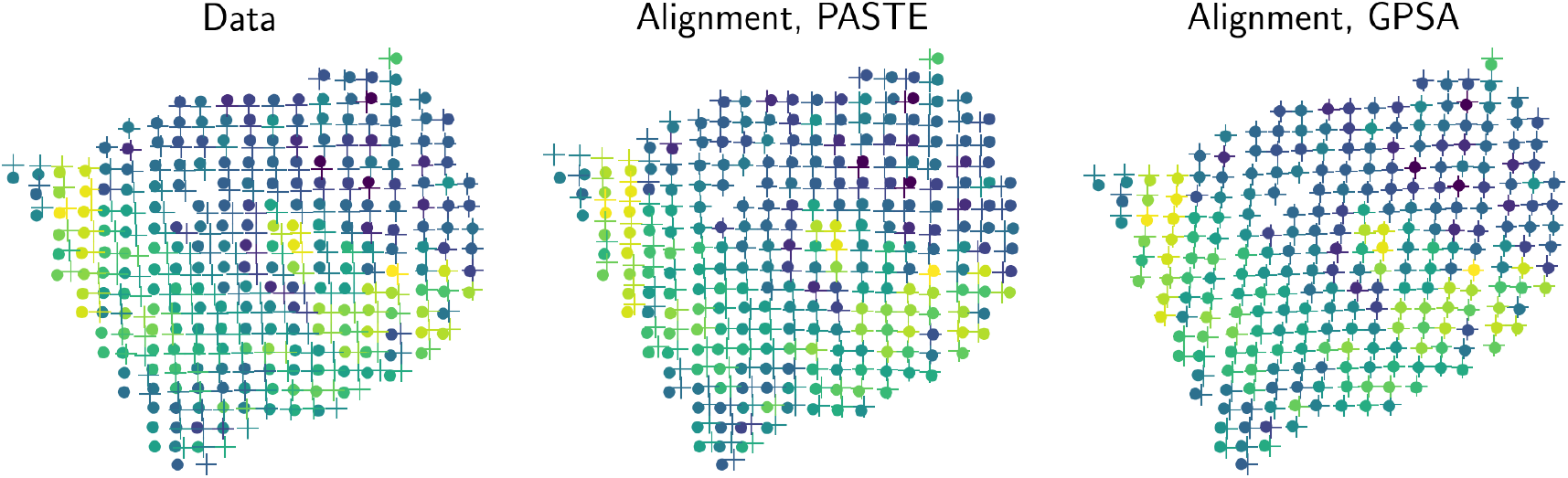
Aligning ST data from a breast cancer tumor. (a) We applied a synthetic warp to one slice of the ST data Ståhl et al. (2016) and sought to remove the resulting spatial distortion. The original slice is plotted using dots, and the synthetic warped slice is plotted using crosses. Points are colored by the expression level of one gene. (b) Alignment from PASTE (Zeira et al., 2021), which applies a linear transformation. (c) Alignment from GPSA.

#### 4.3.2 Estimating expression variability across spatial locations

We asked whether we could estimate the variability of gene expression within each spatial location. An accurate estimate of the variance would allow for useful analyses, such as understanding the spatial heterogeneity of gene expression in a particular tissue region and analyzing changes in expression along the *z* axis. Better alignment of the slices will lead to a more precise quantification of variance.

To quantify expression variation across slices, we again leveraged the ST breast cancer data (Ståhl et al., 2016). Here, we aligned all four slices using a template-based alignment, using the second slice as the template, and assumed they all shared a *z*-axis location as biological replicates. We then transferred each slice’s observed gene expression onto the template slice’s spatial coordinates by assigning each point in the template slice the average of its nearest neighbors in the corresponding aligned slice. Using these four slices within the common coordinate system, we then computed the variance for each gene (Figure 6a) representing a combination of experimental, biological, and *z*-axis variability.

**Figure 6:**
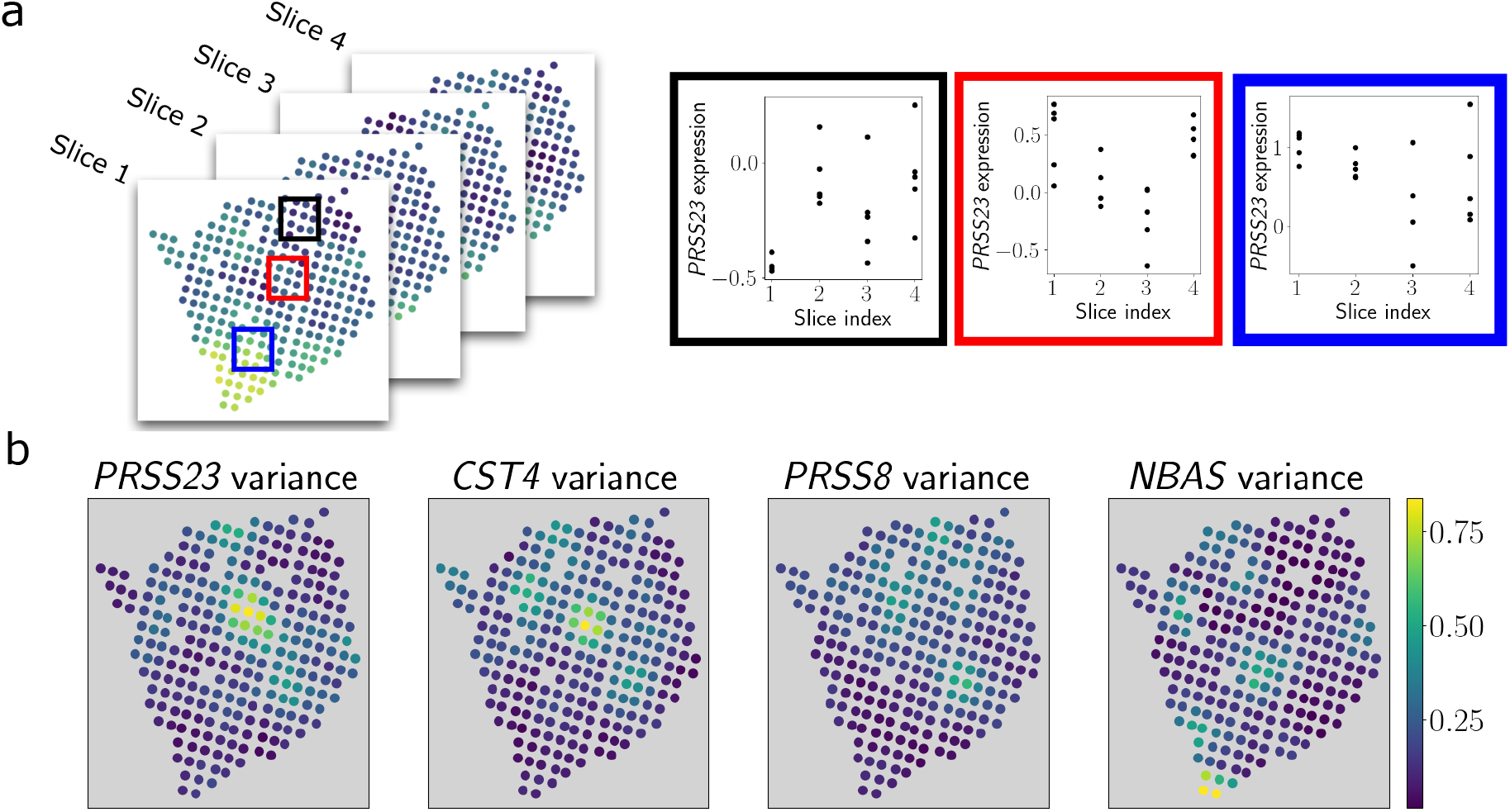
Estimating gene expression variability using GPSA alignments. (a) Using the aligned spatial coordinates provided by GPSA of the four ST slices, we estimated the variability of gene expression within each spatial location by computing the variance across slices for each gene. (b) Several genes show substantial variability across the slices. Points are colored by the estimated variance at each spatial location.

We found substantial variability in gene expression levels across the tissue for several genes (Figure 6b). Moreover, several of these genes have been shown to be related to tumor progression, including *PRSS23* (Chan et al., 2012) and *CST4* (Zhang et al., 2017). These results suggest that studying the variability of gene expression across aligned slices could allow the exploration of spatial gene expression variation at single locations.

#### 4.3.3 Aligning samples in three-dimensional space

Our analyses of the ST data so far have ignored the three-dimensional nature of the contiguous slices. Thus, we next asked whether we could include information about the third spatial dimension (the *z* axis) to create a common coordinate system plus localized gene expression distributions for the three-dimensional tumor — what we would call a *tumor atlas* (Rozenblatt-Rosen et al., 2020).

*To create a three-dimensional tumor atlas, we fit GPSA on the four ST breast cancer slices, but this time we set the number of spatial dimensions to D* = 3. We initialized the four slices’ *z* axis coordinates as [0, 1, 2, 3] and allowed the model to warp these coordinates so that the spatial relationships of the expression data were consistent across spatial dimensions. Importantly, we used the same covariance function parameters for the warping GP across all spatial dimensions, which allows the alignment along the *z* axis to be informed by spatial relationships along the *x* and *y* axes.

To perform the alignment, we used a two-step procedure. In the first step, we performed a template-based alignment with the second slice as the fixed template. In the second step, we fixed the aligned coordinates from warped slices one, three, and four as the template, and fit GPSA again, allowing the second slice’s coordinates to be adjusted.

This process resulted in a three-dimensional common coordinate system for the breast cancer tumor, where we have an estimate of gene expression distribution at each location in this three-dimensional space. We found that the aligned *z* axis coordinates showed substantial adjustments from their original positions (Figure 7). Because we are using GPs, we have estimates of expression distributions for any location inside the grid or outside, whether that location has been observed or not.

**Figure 7:**
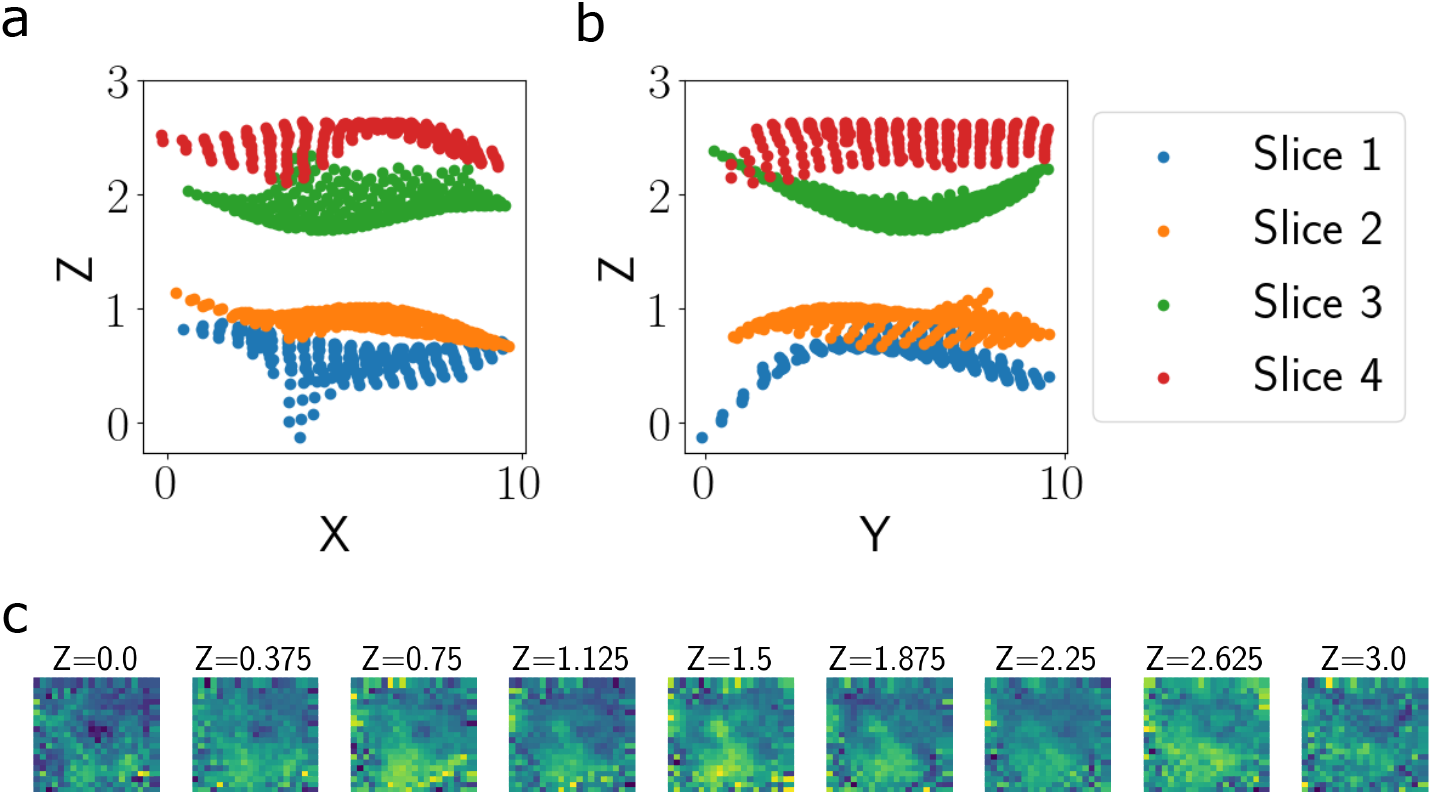
Three-dimensional alignment of ST breast cancer data. Aligned three-dimensional coordinates plotted from two views. (a) shows the *x* axis vs. the *z* axis, and (b) shows the *z* axis vs. the *y* axis. Each panel in (c) shows the imputed gene expression values for the gene *FN1* in one slice of the *z* axis. The location along the *z* axis increases left to right and top to bottom.

Moreover, we used our three-dimensional probabilistic tumor atlas to explore the spatial organization of gene expression within the tumor. We imputed a dense three-dimensional model of gene expression within the learned common coordinate system (Supplementary Fig 2). We found that expression levels of specific genes varied smoothly across this space, with substantial variation along the *z* axis (Figure 7). These findings suggest that GPSA is a feasible model for creating atlases using spatial genomics slices from a single tissue sample.

#### 4.3.4 Aligning Visium profiles of mouse cortex

Next, we applied GPSA to data collected using the Visium platform developed by 10x Genomics (10x Genomics, 2020). These data — which contain two adjacent slices — were collected from a cross section of the sagittal-posterior region of a mouse brain. The two slices contain measurements at 3, 355 and 3, 289 spatial locations, respectively. We again filter the datasets to keep genes that are spatially variable using a nearest-neighbor approach (Appendix 6.3.8), leaving us with 135 genes.

We fit template-based GPSA, designating the first sample as the template, to these samples in order to align them to a shared coordinate system and remove distortions that appeared during tissue collection or sequencing. We then extracted the aligned coordinates. In the original data, there was a small spatial mismatch in the cerebellar folds of the two slices (Figure 8). Examining the aligned coordinates, we found that GPSA was able to correct this distortion by adjusting the second slice downward to match the first slice (Figure 8).

**Figure 8:**
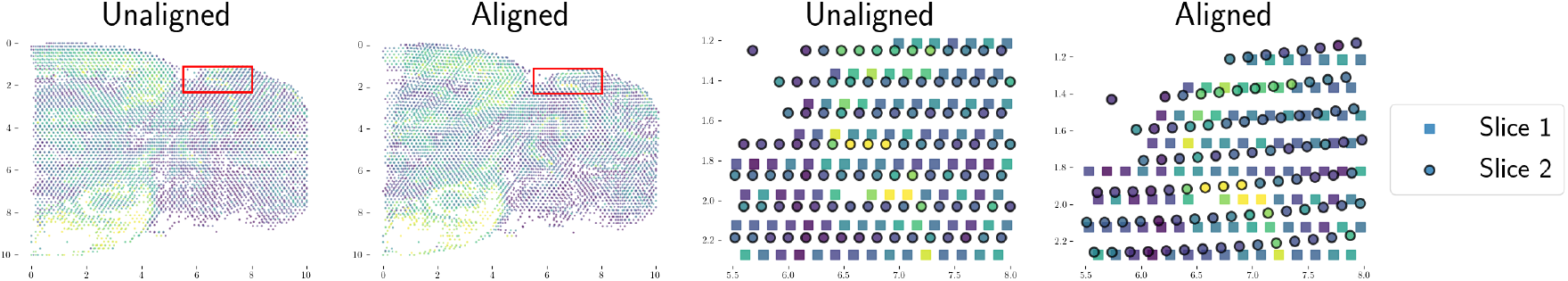
Aligning serial spatial gene expression slices from mouse cortex from the Visium platform. We applied GPSA to two adjacent slices collected from the posterior sagittal region of the mouse brain. The top row shows a superposition of the two slices in the unaligned coordinate system (top left) and the aligned coordinate system (top right). The bottom row shows a magnified version of the region in the red bounding box in the top row. We observe that, in the original slices, a cerebellar fold is misaligned (bottom left). We find that the GPSA common coordinate system corrects this distortion (bottom right).

To quantitatively measure the quality of the estimated alignment, we performed a prediction experiment similar to the one performed using simulated data in subsubsection 4.2.3. We fit GPSA using all of the spots from the first slice and 80% of the spots from the second slice, reserving the remaining 20% of the spots for testing. We then made predictions for the gene expression levels at the held-out spots using two strategies: 1) Naively stacking the two slices and making predictions using a GP (the union GP), and 2) Using our estimated common coordinate system and localized expression distributions to predict the spot expression values. We computed the *R*^2^ for these predictions, repeating this experiment five times for random train/test splits. We found that predictions using the aligned coordinates from GPSA slightly outperformed those using the original coordinates (Figure 9). This finding implies that GPSA is estimating a coordinate system that captures functional similarity and removes distortion between slices.

**Figure 9:**
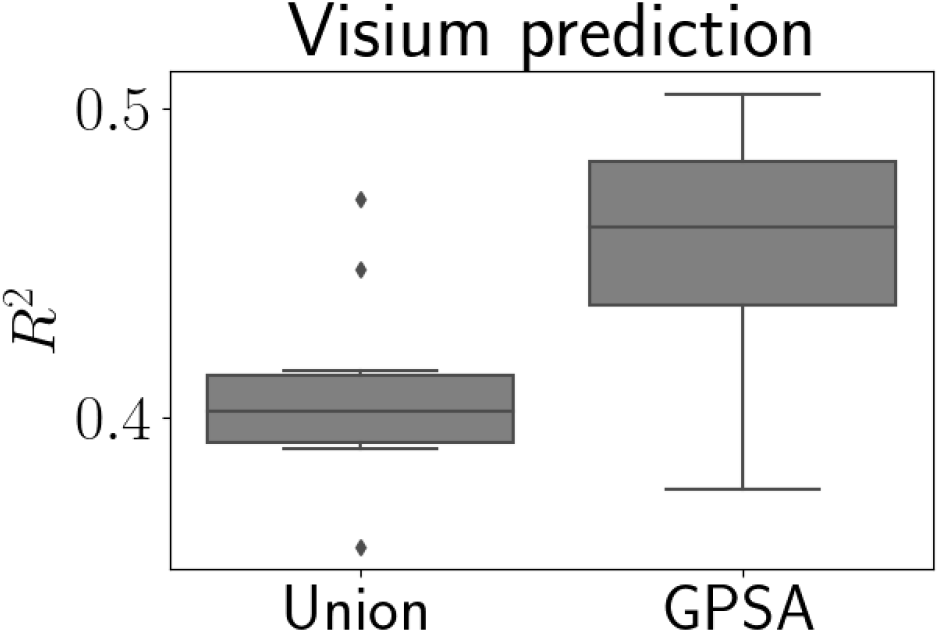
Prediction experiment with Visium data. Coefficient of determination (*R*^2^) for predictions of readout values versus ground truth on a held-out dataset. “Union” represents predictions from a GP fit to a naive concatenation of the observed samples, and “GPSA” refers to predictions from a GP fit jointly across all samples using the aligned coordinate system.

#### 4.3.5 Aligning Slide-seqV2 profiles of the mouse hippocampus

We next leveraged a set of two tissue slices collected from the hippocampus region of two mice using Slide-seqV2 (Stickels et al., 2021). Compared to the Visium platform, Slide-seqV2 has a much higher spatial resolution, capturing RNA levels at near-cellular resolution.

These samples are not immediately comparable due to major shifts in the field-of-view (Supplementary Fig 3). Thus, as a preprocessing step, we first applied a coarse manual rotation and translation to put the samples approximately within the same field-of-view. We fit GPSA to these slices using a template-based alignment with the first slice chosen as the template. We then examined the resulting aligned samples.

We found that the aligned coordinates showed correspondence between the two slices for multiple major landmark regions. In particular, we found that the dentate gyrus and CA1-3 pyramidal layer (also known as Ammon’s horn) from the two slices were well aligned (Figure 10). Due to differences in the field-of-view between the two slices and in the brains of two different mice, we do not expect to achieve a perfect one-to-one matching of the spatial coordinates. In particular, we observe that the choroid plexus is a prominent marker in the first slice but not the second (Supplementary Fig 4), as well as several other structures that are not present in the second slice (Supplementary Fig 5).

**Figure 10:**
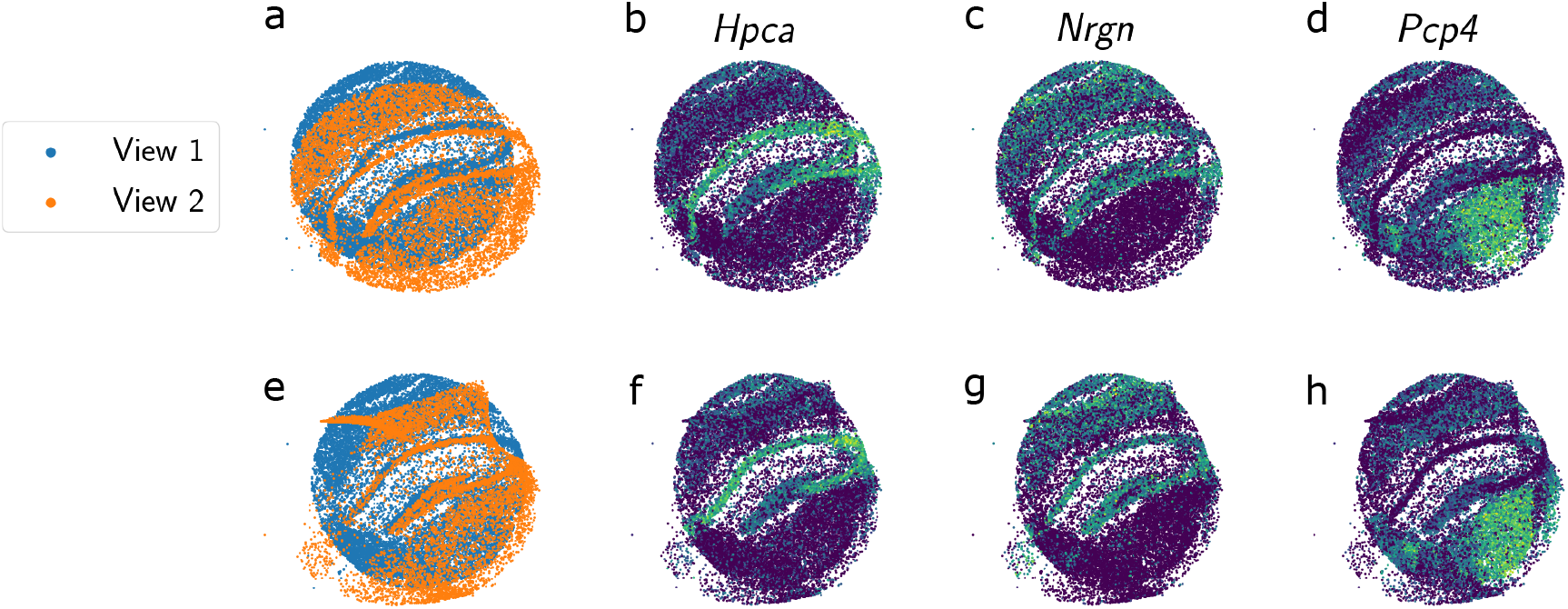
Alignment of Slide-seqV2 spatial gene expression profiles collected from the mouse hippocampus. Each plot in the top row shows a superposition of the two unaligned slices, and each plot in the bottom row shows a superposition of the aligned slices. (a) and (e) are colored by the slice identity, and (b)-(d) and (f)-(g) are colored by the expression levels of single genes (b,f: *Hpca*; c,g: *Nrgn*; d,h: *Pcp4*).

### 4.4 Multi-modal alignment: Incorporating histology images

Finally, we jointly aligned spatially-resolved gene expression and histology images. While the histology images are much lower-dimensional in terms of the outcome variable, which contains measurements for three image channels, these images often contain salient and interpretable features of anatomy and structure. Furthermore, these images are widely used by pathologists, leading to their availability alongside spatial gene expression profiling. We hypothesize that including histology images in the alignment procedure may produce better alignments and further enhance the interpretability of our alignments in downstream analyses, such as enabling annotation of tissue structures in the common coordinate framework. To test this hypothesis, we again leveraged the mouse brain data from the Visium platform. For each of the slices, a histology image is also available and pre-aligned to the corresponding gene expression spatial locations. We again fit template-based GPSA using the *S* = 2 slices and designating the first slice as the template, but this time we also include the histology images as phenotypic readout features and spatial locations. In this setting, GPSA encourages spatial locations with similar patterns in both gene expression and histology to be aligned. We extract the aligned coordinates from the model.

We found that the model successfully aligns these multi-modal samples (Figure 11). We observed that, in the original histology stains, there was a slight mismatch between the slices in one of the cerebellar folds (Figure 11c). After fitting GPSA, we observed that the fold had been corrected in the aligned coordinate system (Figure 11d). Examining the corresponding region of gene expression, we found that certain expression patterns also localized in this area. For instance, the darker histology region corresponded to higher levels of expression in the genes *CAMK2A* (Benjamini Hochberg adjusted p-value *<* 10^*−*5^) and *MT-CO1* (adjusted p-value = 0.005) (Supplementary Fig 6).

**Figure 11:**
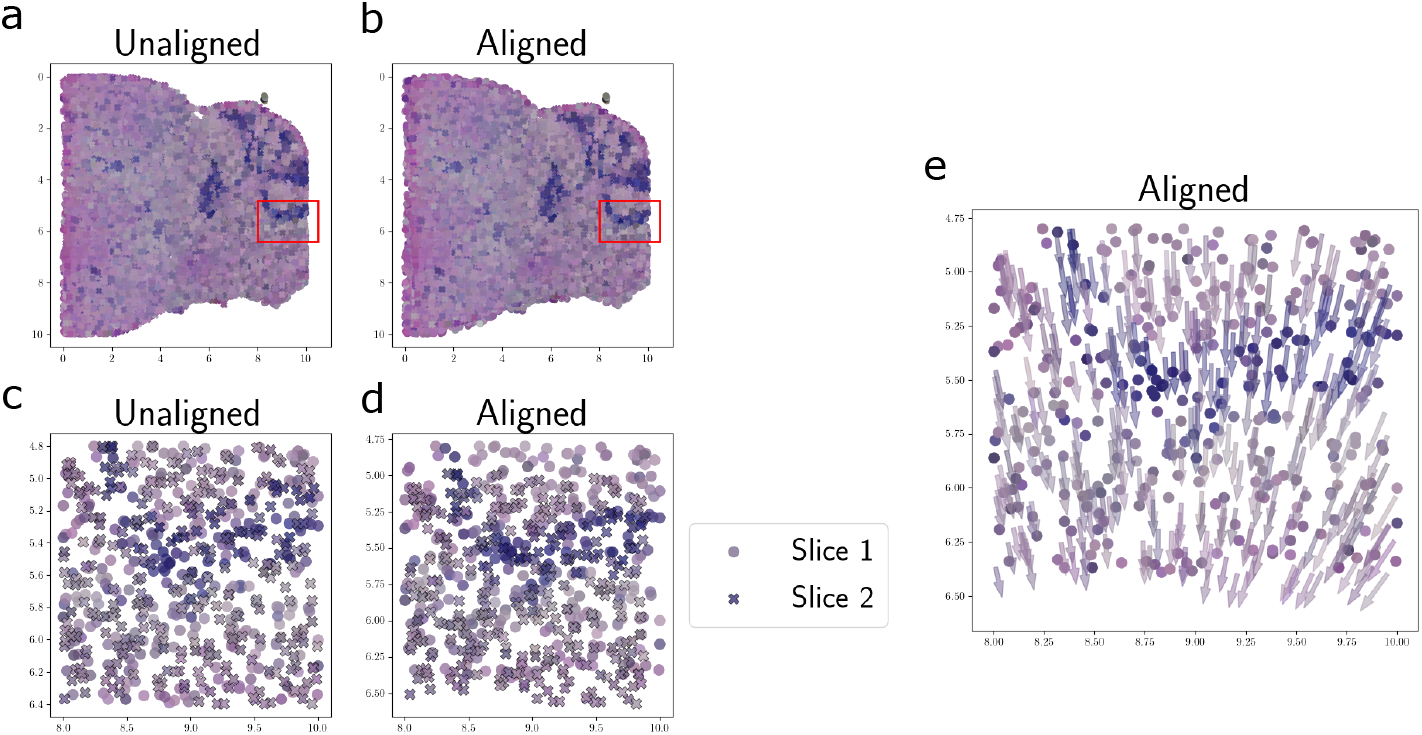
Joint alignment of spatial gene expression and histology images using mouse brain data collected with the Slide-seqV2 platform. (a) and (b) show a superposition of the histology images of the two slices in the unaligned and aligned coordinate systems, respectively. (c) and (d) show magnified versions of the region in the red bounding box. (e) shows the same data as (d), but the points for the second slice are shown with arrows pointing in the direction of the alignment.

To more closely examine GPSA’s behavior in this setting, we computed a vector field demonstrating the displacement of each spatial coordinate after the warp. We observed that substantial nonlinear warping was necessary in order to align the histology stains (Figure 11e). These results suggest that GPSA may feasibly be used to align multi-modal data including spatial gene expression and histology measurements, broadening its potential applications.

## 5 Discussion

We have presented Gaussian Process Spatial Alignment (GPSA), a Bayesian model for aligning multiple spatial genomics and histology slices in a known or unknown common coordinate system. We have shown that our model can flexibly align samples from multiple spatial sequencing technologies, fields-of-view, and data modalities.

Previous approaches for alignment of spatial genomics slices, such as PASTE (Zeira et al., 2021) and Splotch (Äijö et al., 2019), have relied on linear transformations of the observed spatial coordinates. However, linear transformations do not account for local nonlinear distortions in the coordinate systems. In this paper, we demonstrate the necessity of allowing these finer-scale and nonlinear adjustments in a variety of data types. GPSA models local, nonlinear transformations of spatial coordinates, which flexibly capture many types of warps but require that the warps are fairly small. We could imagine a strategy for building tissue atlases that consists of two stages: 1) Running PASTE to obtain an initial coarse alignment, and 2) Running GPSA to tune the local alignment to a coordinate system and produce a common coordinate system for the slices with localized measurement distributions across the space.

We applied GPSA to data from three spatial gene expression technologies: Spatial Transcriptomics (Ståhl et al., 2016), Visium (10x Genomics, 2020), and Slide-seqV2 (Stickels et al., 2021), plus pre-aligned histology images. Given the flexible nature of GPSA’s modeling assumptions, our model is applicable to many other spatial sequencing technologies (both present and future) with varying levels of resolution, fields-of-view, and depth of phenotypic profiling.

Several future directions could be pursued. The computational complexity of GPSA can be a burden for certain datasets. While we observe that GPSA obtains more accurate alignments compared to competing parametric approaches, such as PASTE (Zeira et al., 2021), this comes at the cost of time. Our variational inducing point approach to inference reduces the time complexity from *O*(*n*^3^) to *O*(*nm*^2^), where *n* and *m* are the number of data points and inducing points, respectively. However, a large number of inducing points are often required for high-resolution spatial genomics technologies, which can result in our approximation being too slow to fit. Nonetheless, we find that the times required to fit GPSA to the data presented here are not prohibitive (Supplementary Fig 7), since researchers only need to align their samples once with GPSA, and any downstream analysis on the aligned samples will not require a realignment. Including known anatomical landmarks in the model could speed up the alignment and lead to more biologically faithful and interpretable coordinate systems. However, we also note that an attractive feature of GPSA is its reliance on almost no prior knowledge about the structure of the tissue. Future work may also use landmarks inferred from the histology images to annotate the inferred shared coordinate system to automate this process.

Additionally, refining the process of filtering phenotypic readout features (e.g., genes) with no spatial variability could further improve the speed and accuracy of our inference approach, as GPSA exploits spatially-variable patterns to perform the alignment. While we have proposed a simple nearest-neighbor approach for filtering features in this work, further work could be done to improve this step. Additional study to determine when it is most appropriate to use a low-rank model of the readout features (as opposed to modeling each feature independently) in the context of alignment could be warranted. The linear model of coregionalization model could also be extended across data modalities to explicitly capture the relationship across data modality features.

Finally, there remains an opportunity for a deeper theoretical study of GPSA and DGPs in general. Studying the posterior consistency of the kernel parameters and the latent variable **G** could lead to theoretical guarantees for the resulting common coordinate systems.

## Author’s contributions

AJ, FWT, DL, and BEE designed the method. AJ implemented the method and conducted data analysis. AJ, FWT, and BEE analyzed the results. AJ wrote the manuscript. AJ, DL, FWT, and BEE edited the manuscript.

## Acknowledgements

This work was funded by Helmsley Trust grant AWD1006624, NIH NCI 5U2CCA233195, NIH NHLBI R01 HL133218, and NSF CAREER AWD1005627.

## 6 Appendix

### 6.1 Data acquisition

#### 6.1.1 Spatial Transcriptomics

ST data were obtained from the PASTE code repository: https://github.com/raphael-group/paste. All four layers from the sample data/ directory were used.

#### 6.1.2 Visium

Visium data were obtained from the 10x Genomics website. Data for the two slices were downloaded from the “Datasets” page. Specifically, spatial gene expression and H&E stains were downloaded from the following links:

- Mouse Brain Serial Section 1 (Sagittal-Posterior): https://www.10xgenomics.com/resources/datasets/mouse-brain-serial-section-1-sagittal-posterior-1-standard-1-1-0
- Mouse Brain Serial Section 2 (Sagittal-Posterior): https://www.10xgenomics.com/resources/datasets/mouse-brain-serial-section-2-sagittal-posterior-1-standard-1-1-0

#### 6.1.3 Slide-seqV2

Slide-seqV2 data were downloaded from the Broad Institute’s Single Cell Portal: https://singlecell.broadinstitute.org/single_cell/study/SCP815/highly-sensitive-spatial-transcriptom Two pucks corresponding to the mouse hippocampus were used: Puck_191204_01 and Puck_200115_08.

### 6.2 Code

Code for the model and experiments is available at https://github.com/andrewcharlesjones/spatial-alignment.

### 6.3 Methods

#### 6.3.1 Gaussian processes

A Gaussian process (GP) is a stochastic process defined as a collection of random variables where any subset follows a multivariate Gaussian distribution. Specifically, *y*_1_, *y*_2_, … constitute a GP if, for any finite set of indices *i*_1_, *i*_2_, …, *i*_*n*_, it holds that

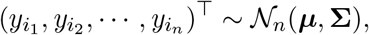

where ***μ*** is a mean vector and **Σ** is a positive definite covariance matrix. GPs are widely used in functional data analysis, machine learning, and spatial statistics due to their flexibility and expressiveness in modeling complex dependent data (Rasmussen and Williams, 2005; Stein, 1999; Gelfand et al., 2010; Cressie and Wikle, 2011; Banerjee et al., 2014). For example, in nonparametric regression models, GPs are commonly used to model unknown arbitrary functions; in Bayesian contexts, they act as priors over functions (Ghosal and Van der Vaart, 2017).

GPs are often used as prior distributions over functions, as in this paper. In this case, for a function *f* defined on the domain ℝ^*D*^, we denote a GP prior as

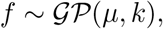

where *μ* : ℝ^*D*^ → ℝ is a mean function and *k* : ℝ^*D*^ × ℝ^*D*^ → R is a positive definite covariance function (also known as a kernel function or covariogram). For noisy responses from the noiseless function *f*, we include Gaussian noise: *y* ∼ 𝒩 (*f* (**x**), *∼*^2^), where *∼*^2^ is often referred to as the *nugget*.

#### 6.3.2 Deep Gaussian processes

Deep GPs (DGPs) were developed to further extend the expressivity of GPs (Damianou and Lawrence, 2013; Salimbeni and Deisenroth, 2017). DGPs are a composition of functions, each of which is drawn from a GP. In the univariate case, the a function drawn from an *L*-layer DGP is given by

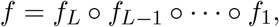

where for each ℓ = 1, …, *L*, we have *f*_ℓ_ ∼ 𝒢 𝒫 (*μ*_ℓ_, *k*_ℓ_), and **y**_ℓ_ = *f*_ℓ_ (*f*_ℓ*−*1_(° ° ° *f*_1_(**x**))) is the output of the ℓth layer an input sample **x**. In this work, we use two-layer DGPs, or *L* = 2.

#### 6.3.3 Stochastic variational inference for GPSA

Although closed-form posterior distributions are available in GPs, this is not the case in DGPs. To perform approximate inference, we leverage a sparse GP framework using inducing points (Titsias, 2009; Snelson and Ghahramani, 2006; Salimbeni and Deisenroth, 2017). Because GPSA is a two-layer DGP, we include inducing points at each of the two layers. In particular, suppose we have a set of *M*^*s*^ *< n*_*s*_ inducing locations (also known as pseudo-inputs) for each slice 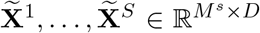, and another set of *M < N* inducing locations in the common coordinate system layer 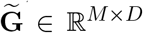. We then denote the associated set of inducing values (pseudo-outputs) for the two layers as 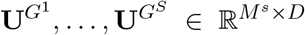 and **U**^*F*^ ∈ ℝ^*M ×p*^, respectively. The joint model (omitting the dependence on the covariance function parameters Θ) is then

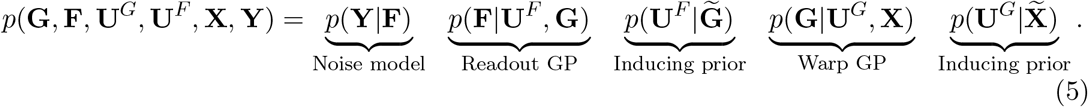

Note that *p*(**F**|**U**^*F*^, **G**) and *p*(**G**|**U**^*G*^, **X**) have closed forms because they are conditional multivariate Gaussians. If a Gaussian noise model is assumed, then *p*(**Y**|**F**) also has a closed form. However, inference in this model scales cubically with the number of spots, so we seek a faster variational approach.

We now specify a variational model *Q*, whose parameters we will optimize to approximate the exact posterior (Equation 2). Following earlier work (Damianou and Lawrence, 2013), we use the following form for the approximate posterior:

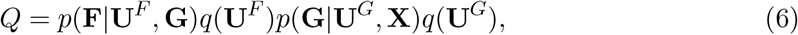

where *q*(**U**^*F*^) and *q*(**U**^*G*^) are chosen to be multivariate normal distributions. We denote the variational parameters collectively as *ϕ*. Because all distributions are Gaussian, we can analytically marginalize out the pseudo-outputs **U**^*F*^ and **U**^*F*^ (Salimbeni and Deisenroth, 2017). See Appendix 6.3.4 for details.

The optimization problem is then to minimize the KL divergence from the exact posterior (Equation 2) to the approximate posterior (Equation 6) with respect to the variational parameters. This is equivalent to maximizing a lower bound on the log marginal likelihood ℒ ≤ log *p*(**Y**) (the evidence lower bound, or ELBO). The variational parameters *ϕ* are made up of the parameters of the prior distributions for the pseudo-outputs, *q*(**U**^*F*^) and *q*(**U**^*G*^), and optionally the inducing locations 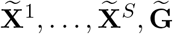. More precisely, our optimization problem is

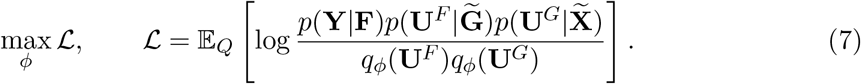

We provide a complete derivation and explanation of this lower bound in the next section.Although this lower bound cannot be evaluated in closed form, we can efficiently sample from it and use these samples to maximize with respect to the variational parameters *ϕ*.

#### 6.3.4 Maximizing the ELBO in GPSA

Recall that the evidence lower bound for a generic model with observed data *x*, latent variable *z*, and approximating distribution *q* is given by

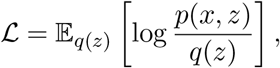

where *p*(*x, z*) is the joint model density, and *q*(*z*) is the variational distribution.

Plugging in our GPSA model, the ELBO is given by

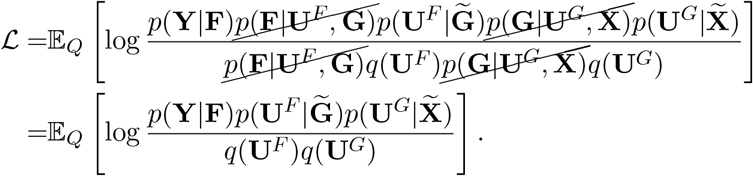

We can split Equation 7 into a term containing the expected log likelihood and two terms that are KL divergences:

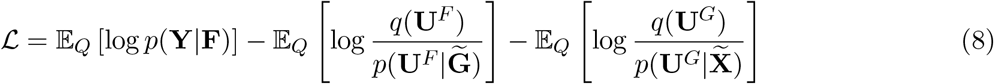

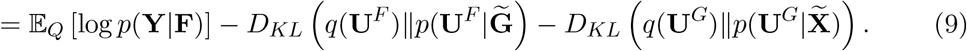

Because we let *q*(**U**^*F*^) and *q*(**U**^*G*^) be multivariate Gaussians, the KL divergence terms have closed forms, and only remaining term to estimate is the expected log likelihood (the first term in Equation 9). We estimate this term with a Monte Carlo approximation. Given *T* samples of **F**, our estimate is

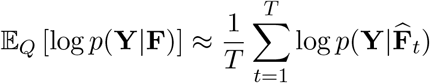

Where 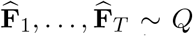. We use a two-staJge procedure to obtain these samples. First, we draw samples of Ĝ from 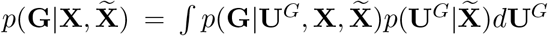. Second, we draw,samples of 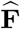 from 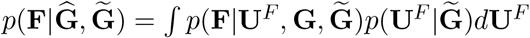. We can write each of these distributions in closed form, which we show next.

Let 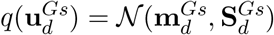. The marginal for **G** is given by

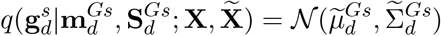

with

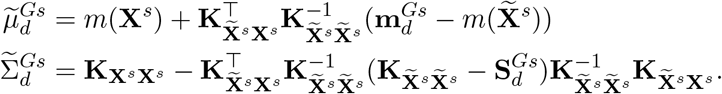

Let 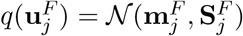. The marginal for **F** is given by with

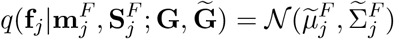

with

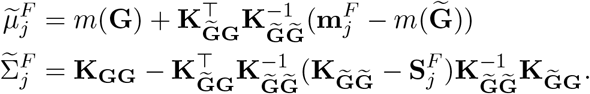

#### 6.3.5 Multivariate correlated outcomes

In its simplest form, GPSA assumes that feature readouts are independent of one another by modeling each with a separate GP-distributed function *f*_*j*_. However, given that our phenotypic readouts of interest (gene expression, for example) are often highly correlated between features, we would like to leverage the correlation between readouts to fit *f*. There are several approaches to accounting for this correlation (Boyle and Frean, 2005; Goulard and Voltz, 1992; Gelfand et al., 2004).

We choose to leverage the linear model of coregionalization (LMC) (Goulard and Voltz, 1992). Rather than allowing *p* separate GPs, the LMC assumes there are *L < p* latent Gaussian processes, and that the observed readouts are linear combination of the outputs of these latent GPs. To incorporate this into our registration model, we assume the following model for the second layer GP:

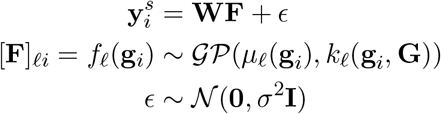

where **W** ∈ ℝ^*p×L*^ is a loadings matrix, and **F ∈ ℝ**^*L×N*^ is a matrix containing latent factors. Given a set of warped coordinates **G**, our likelihood is then

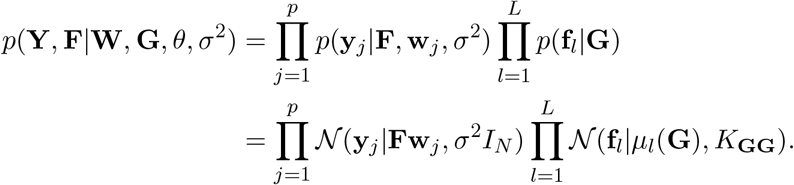

Including the warp model, our entire joint model becomes

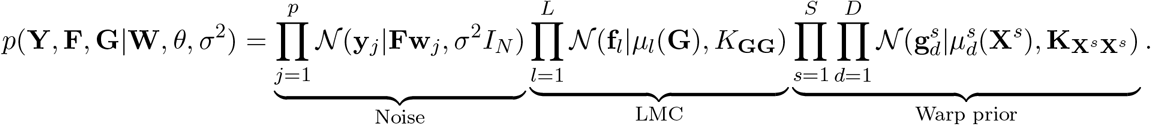

In our applications, we may not be interested in directly estimating the latent factors **F**. We can marginalize these out (Kyzyurova, 2019) and write the likelihood as

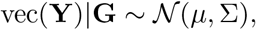

where

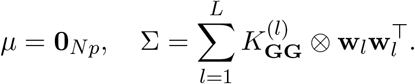

If the latent covariance functions *k*_1_, …, *k*_*L*_ are the same, the covariance simplifies as Σ = *K*_**GG**_ ⊗ **WW**^*T*^.

#### 6.3.6 Multi-modal outcomes

We may sometimes have access to multiple samples from each slice, each of whose phenotypic readouts are collected from different modalities. For example, we may have a spatial transcriptomics sample and a histology image in each slice. While both of these modalities lie in a two-dimensional spatial coordinate system, they have different response values. In this example, the spatial transcriptomic readouts will be **Y**^1^ ∈ ℝ^*n×p*^, where *p* is the number of genes, while the histology image readouts will be **Y**^2^ ∈ ℝ^*m×q*^, where *q* is the number of color channels.

Our model can easily accommodate this setting. We assume that the different modalities are already aligned within each slice, which is a reasonable assumption in practice. Instead of computing the likelihood for just one set of phenotypic readouts, we compute it for each modality’s phenotypic readouts. For example, the likelihood becomes

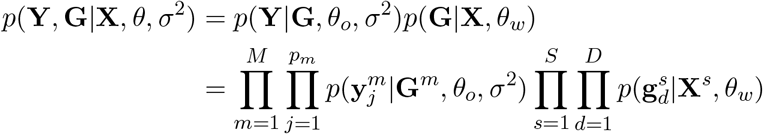

where *M* is the number of modalities, *p*_*m*_ is the number of readout features in modality *m*, and **Y**^*m*^ is the set of readout features for modality *m*.

#### 6.3.7 Non-Gaussian likelihoods

We can easily accommodate non-Gaussian likelihoods in this model. In particular, we can specify the likelihood in Equation 5, *p*(**Y**|**F**) to be any suitable likelihood. In the setting of sequencing data, the measurements often come in the form of nonnegative integer counts, for which a Poisson likelihood is often a reasonable choice.

#### 6.3.8 Nearest neighbors approach to filtering for spatially variable genes

For the spatial gene expression datasets, we use a simple approach to remove genes that do not show noticeable spatial variability. Empirically, we find that removing these genes leads to faster convergence and more accurate common coordinate systems.

To estimate the spatial variability of each gene, we perform a prediction experiment where for each spatial location 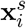, we predict its expression value as the average of its *k* nearest neighbors (we choose *k* = 10). We then compute the coefficient of determination (*R*^2^) between these predictions and the measured expression to estimate the spatial coherence of the expression values. We then choose a threshold *τ* and remove any genes with an *R*^2^ value less than *τ* (we choose *τ* = 0.3).

We find that this simple approach identifies a set of genes with high spatial variability (Supplementary Figures 8, 9) and identifies genes that have been validated by previous work (Svensson et al., 2018). More complicated procedures to identify spatially variable genes could be used (Svensson et al., 2018), but this is not the primary focus of our work.

#### 6.3.9 Synthetic warps

Throughout our experiments, we apply three different types of random warps, which we describe here.

1. *Linear warp*: This warp applies a linear transformation to the observed spatial coordinates for each slice **X**^*s*^ such that 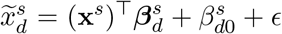 for *d* ∈ {1, …, *D*}, where 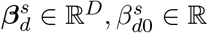, are the slope and intercept, respectively, and ϵ∼ 𝒩 (0, *∼* ^2^) is a noise term.
2. *Polar warp*: For a single spatial sample be represented as **x** = [*x*_1_, *x*_2_]^*T*^, this function is defined as

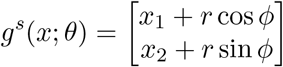

where *θ* = {*r, ϕ*}. We further parametrize *θ* to allow for location-specific distortions. Thus, *θ* is implicitly a function of *x* as well,

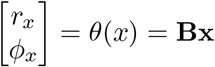

where **B** is a 2 × 2 coefficient matrix. The full warping function can then be written as

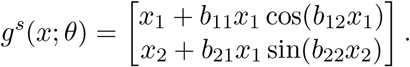
3. *GP warp*: Applies a transformation function that is drawn from a GP:

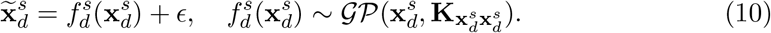

**Supplementary Fig 1:**
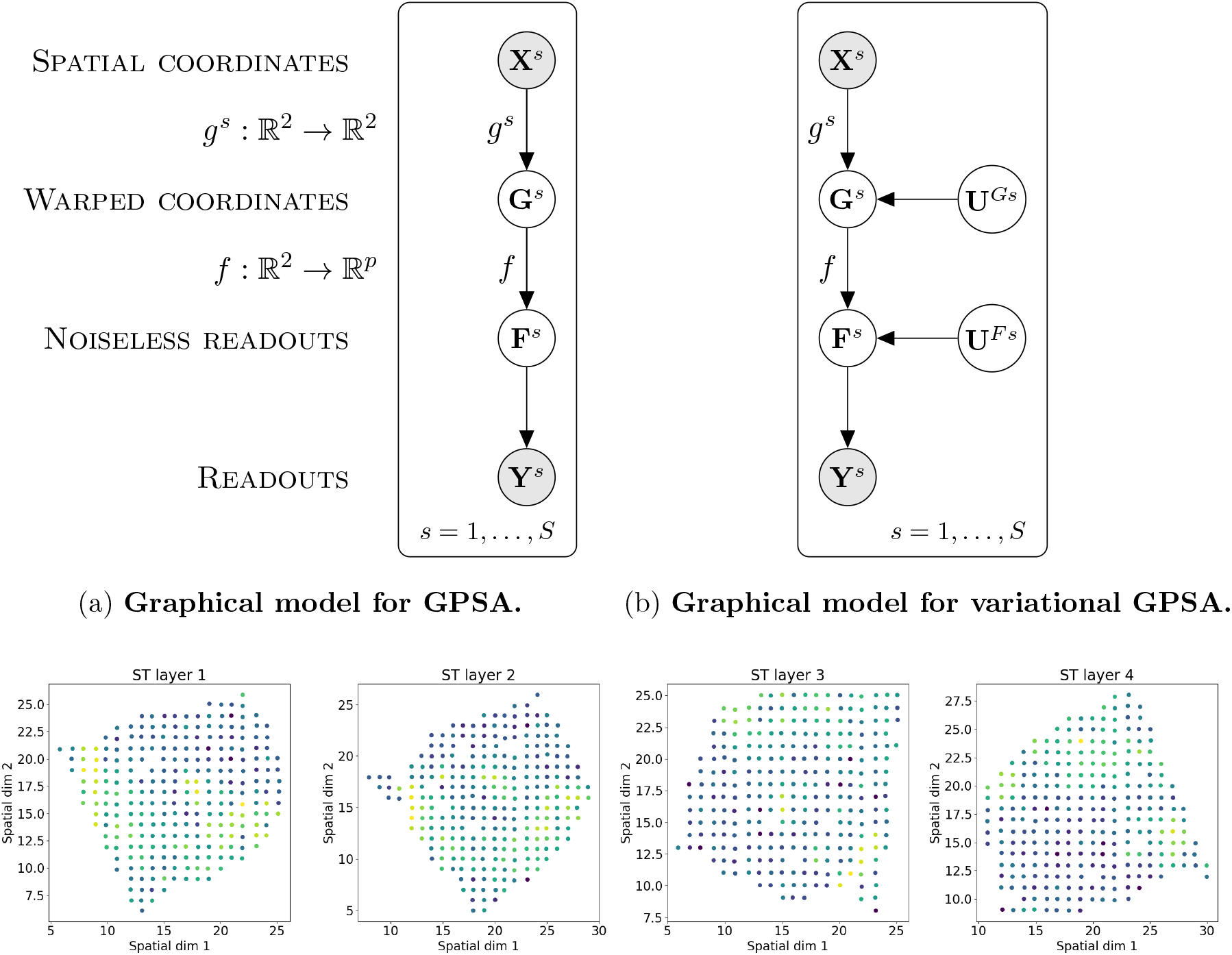
Spatial Transcriptomics sample from breast cancer patient.

## 6.4 Supplementary figures

**Figure 13:**
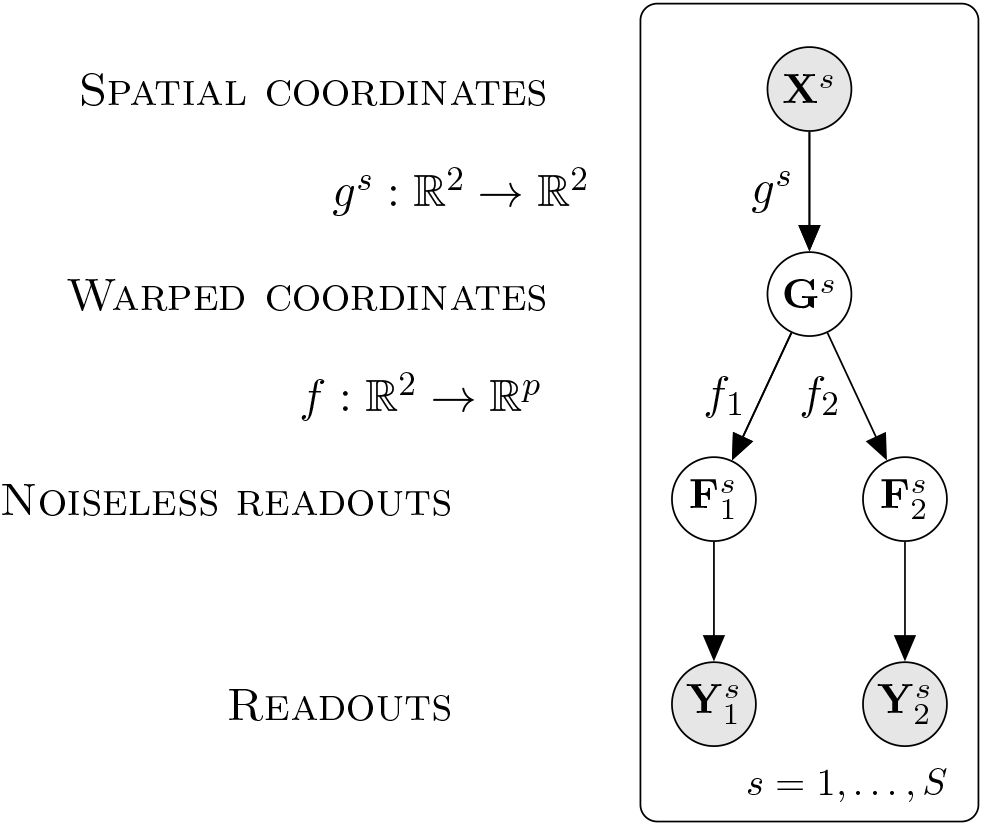
Graphical model for multimodal GPSA.

**Supplementary Fig 2:**
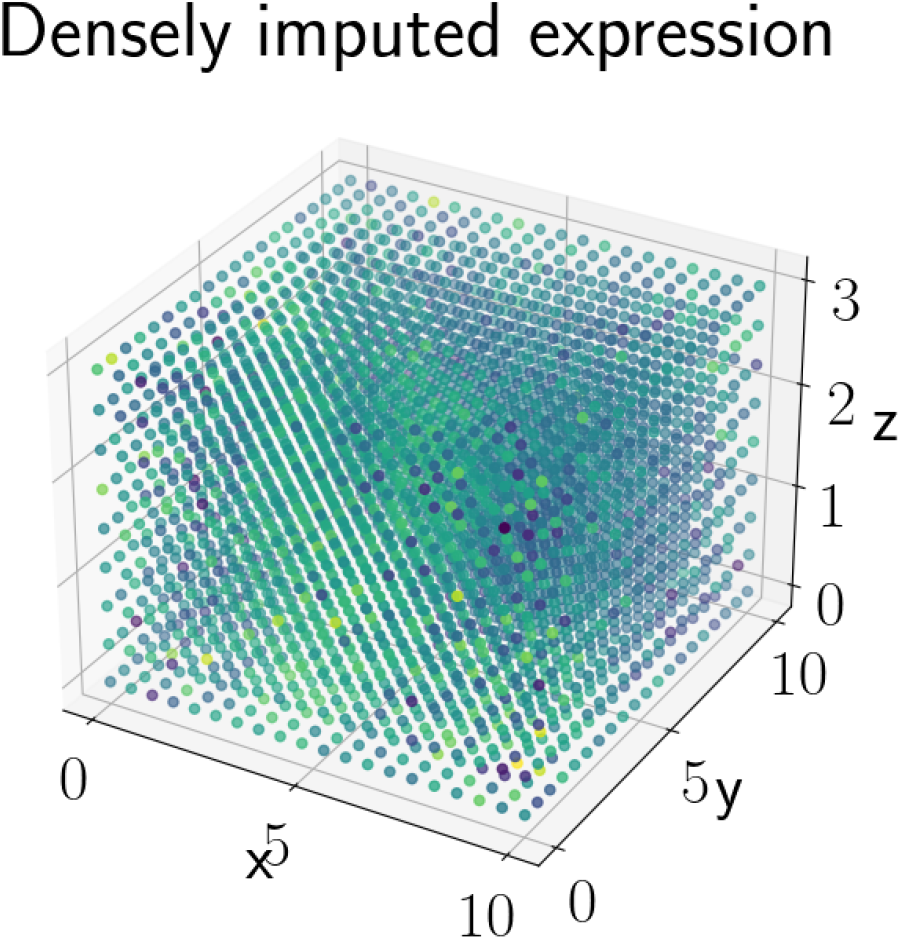
Dense imputation of gene expression within the three-dimensional common coordinate system for the ST breast cancer data.

**Supplementary Fig 3:**
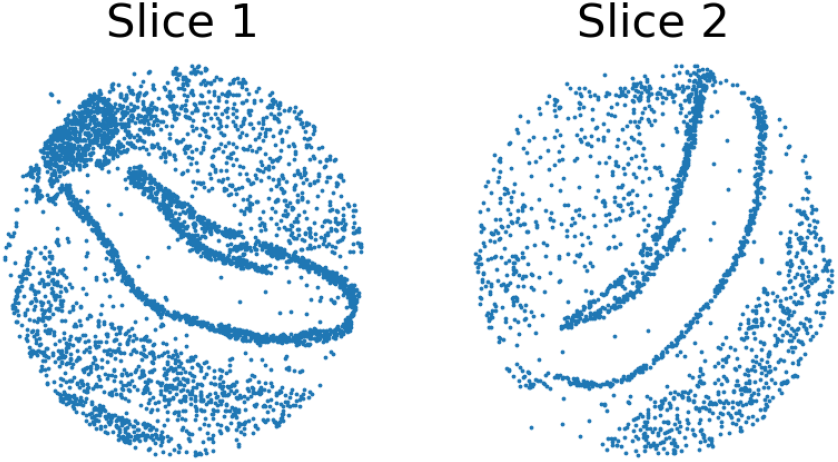
Slide-seqV2 slices without initial coarse alignment.

**Supplementary Fig 4:**
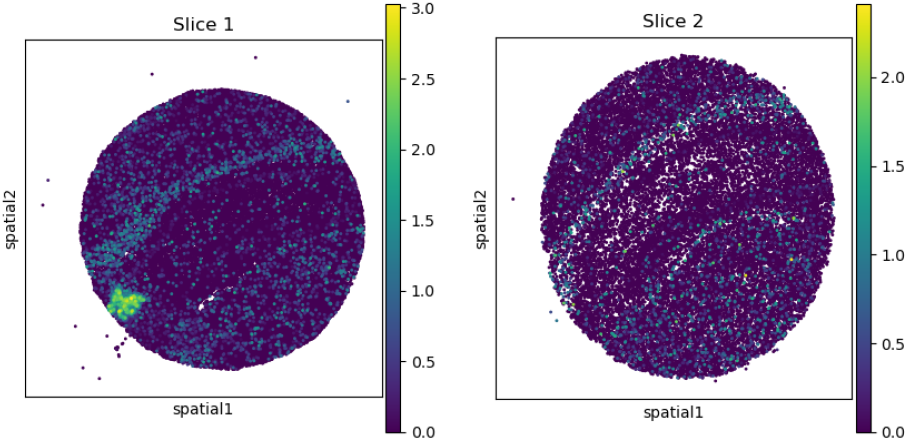
Expression of the gene *ENPP2* in the two Slide-seqV2 slices.

**Supplementary Fig 5:**
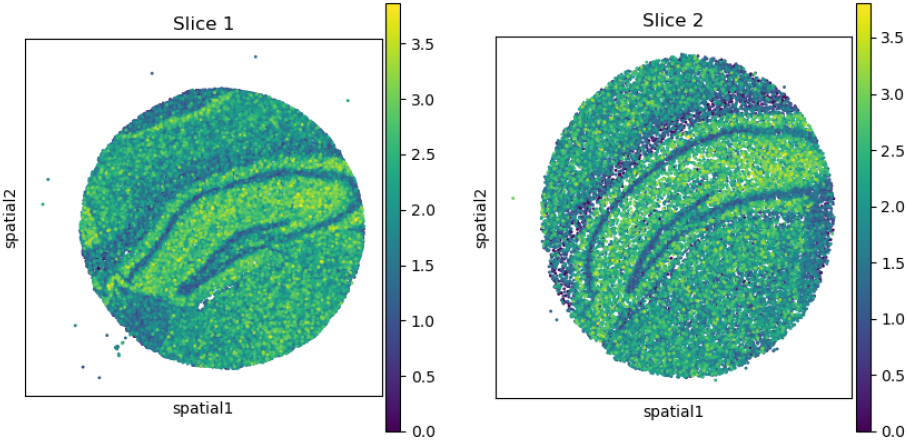
Expression of the gene *MR-ND1* in the two Slide-seqV2 slices.

**Supplementary Fig 6:**
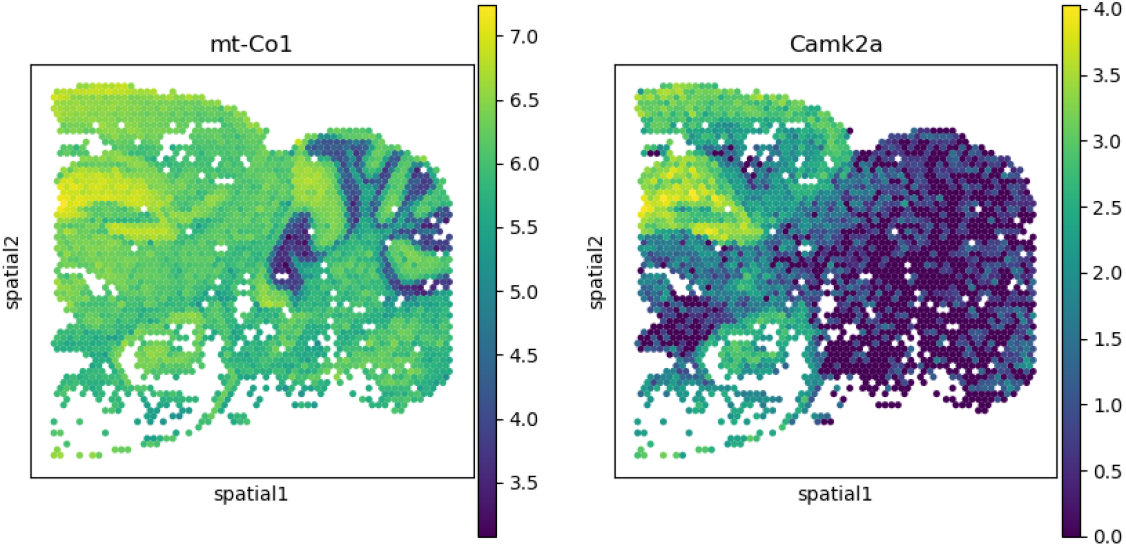
Expression of *MT-CO1* and *CAMK2A* in the Visium mouse cortex.

**Supplementary Fig 7:**
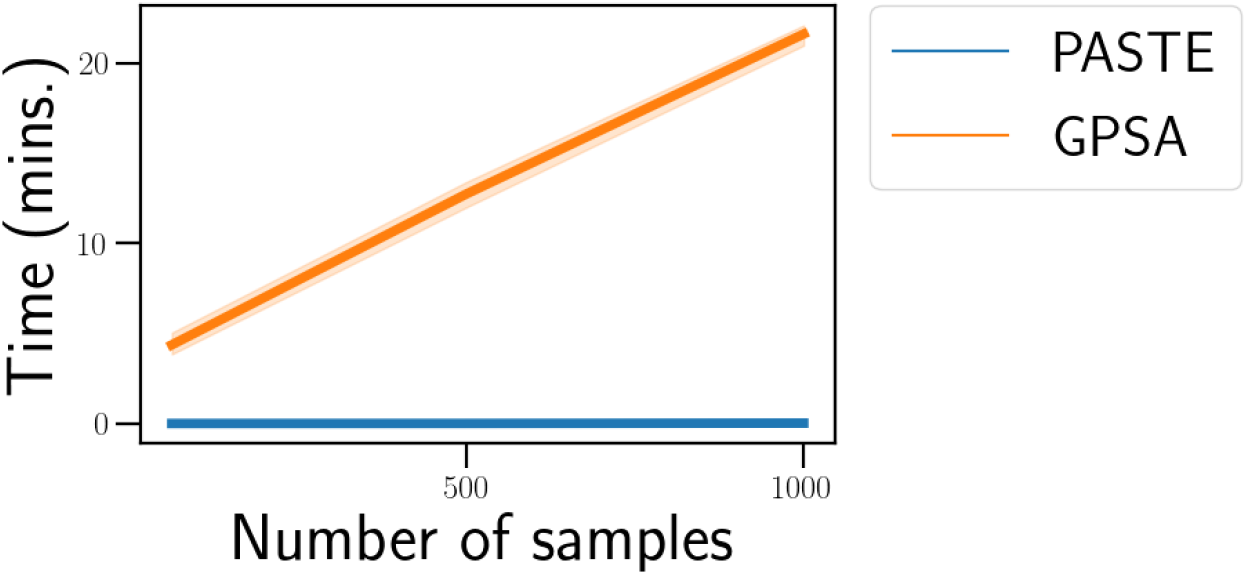
Time complexity of GPSA and PASTE.

**Supplementary Fig 8:**
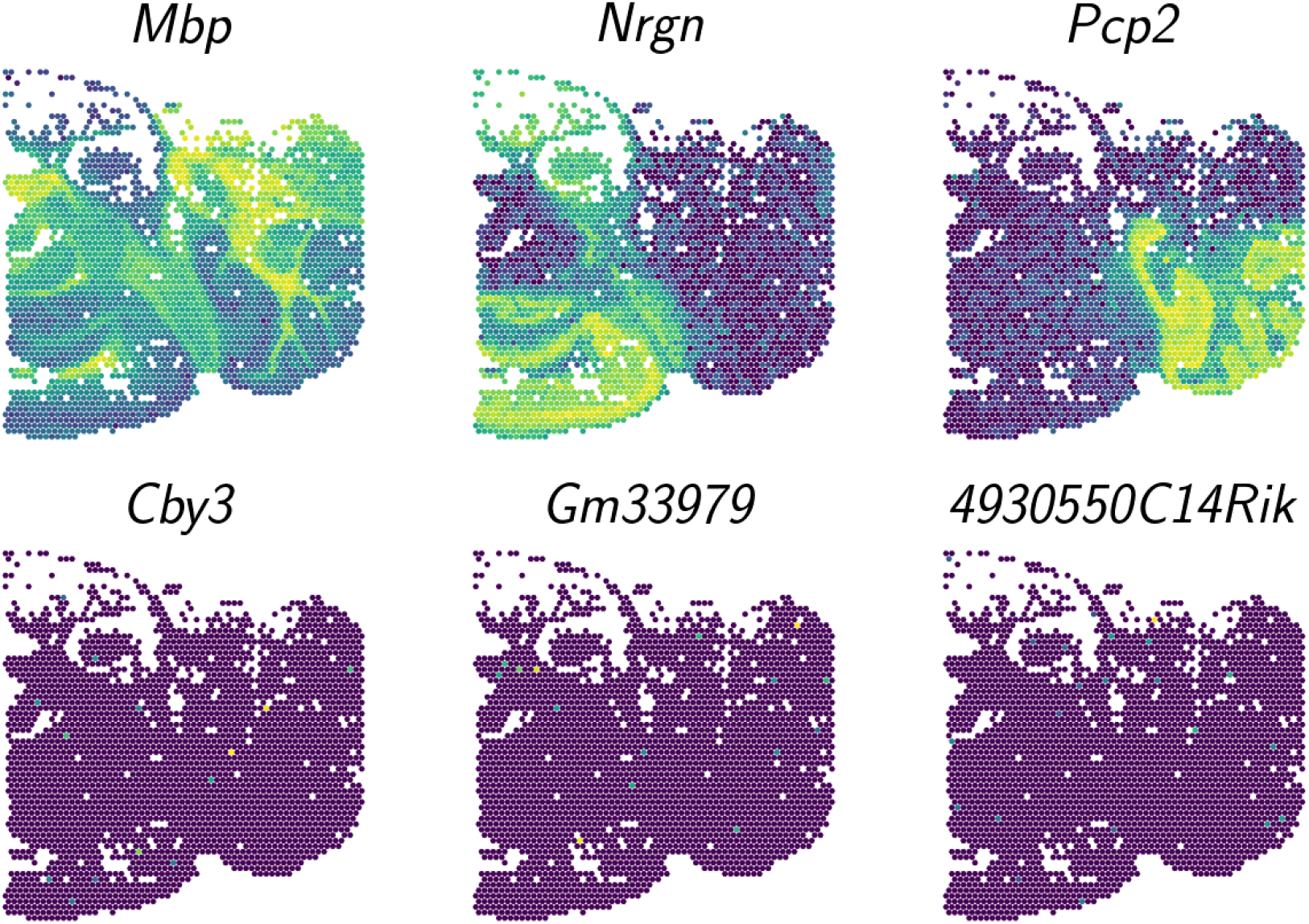
Spatially variable and non-variable genes in the Visium mouse brain dataset. The top row shows the a set of example genes that were spatially variable, and the bottom row shows an example set of genes that were not highly spatially variable.

**Supplementary Fig 9:**
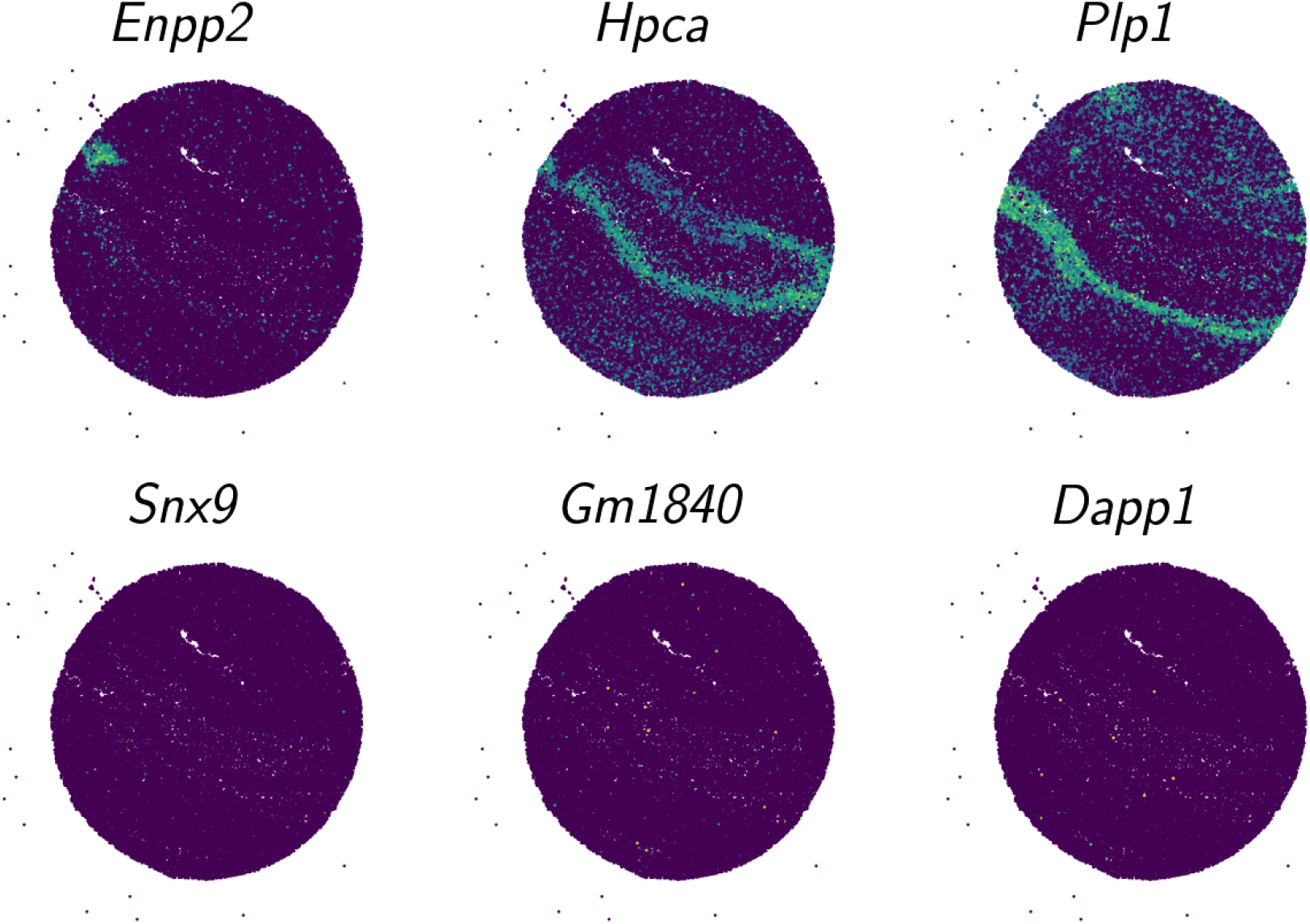
Spatially variable and non-variable genes in the Slide-seqv2 mouse brain dataset. The top row shows the a set of example genes that were spatially variable, and the bottom row shows an example set of genes that were not highly spatially variable.

